# iScore: A ML-Based Scoring Function for *de novo* Drug Discovery

**DOI:** 10.1101/2024.04.02.587723

**Authors:** Sayyed Jalil Mahdizadeh, Leif A. Eriksson

## Abstract

In the quest for accelerating *de novo* drug discovery, the development of efficient and accurate scoring functions represents a fundamental challenge. This study introduces iScore, a novel machine learning (ML)-based scoring function designed to predict the binding affinity of protein-ligand complexes with remarkable speed and precision. Uniquely, iScore circumvents the conventional reliance on explicit knowledge of protein-ligand interactions and full picture of atomic contacts, instead leveraging a set of ligand and binding pocket descriptors to evaluate binding affinity. This approach avoids the inefficient and slow conformational sampling stage, thereby enabling the rapid screening of ultra-huge molecular libraries, a crucial advancement given the practically infinite dimensions of chemical space. iScore was rigorously trained and validated using the PDBbind 2020 refined set, CASF 2016, and CSAR NRC-HiQ Set1/2, employing three distinct ML methodologies: Deep Neural Network (iScore-DNN), Random Forest (iScore-RF), and eXtreme Gradient Boosting (iScore-XGB). A hybrid model, iScore-Hybrid, was subsequently developed to incorporate the strengths of these individual base learners. The hybrid model demonstrated a Pearson correlation coefficient (*R*) of 0.78 and a root mean square error (RMSE) of 1.23 in cross-validation, outperforming the individual base learners and establishing new benchmarks for scoring power (*R* = 0.814, RMSE=1.34), ranking power (*ρ* = 0.705), and screening power (success rate at top 10% = 73.7%).

## 1. Introduction

Molecular docking is undoubtedly the most widely used technique in structure-based computer aided drug discovery that aims to predict the binding mode and binding affinity of small organic molecules toward a target protein.^1^ The performance (speed and accuracy) of a molecular docking program strongly depends on its two main components, sampling and scoring.^2^ Sampling refers to a search algorithm that evaluates a finite number of ligand conformations within and around the binding site of a target protein to elucidate the ligand binding mode. Scoring refers to a class of computational methods, called *scoring functions*, that are formulated to predict the binding affinity of each ligand conformation within the protein binding site.^3^ The performance of a scoring function can be determined by three evaluation metrics:^4^ “*scoring power*” that indicates the degree of correlation in the predicted versus experimentally determined binding affinity values, “*ranking power*” that is the capability of the scoring function to accurately rank a given set of active ligands with respect to their predicted binding affinity values, toward a particular protein target, and “*screening power*” that refers to the ability of the scoring function to identify the true ligand with the highest affinity against a given protein target among a set of random decoy molecules. While the scoring part of a typical molecular docking calculation is relatively fast, the sampling part is time consuming, computationally expensive, and inefficient.^5^ Therefore, in traditional molecular docking and virtual screening, the calculation time and cost scale exponentially with increasing number and degree of freedom of molecules under evaluation. On the other hand, the size of available molecular databases for virtual screening is extremely limited (several million up to a few billion molecules) that covers only a tiny part of the actual chemical space which is predicted to be as large as ∼10^60^ feasible drug-like molecules.^6^

Traditional scoring functions can be categorized into the three main classes force-field based, empirical, and knowledge-based, depending on the way they are formulated.^7^ Despite significant improvements in the last decade, several recent studies clearly show that the performance of traditional scoring functions is quite limited in both scoring power and ranking power aspects.^8^ On the other hand, the most successful scoring approaches such as free energy perturbation (FEP) techniques,^9^ are very sensitive to the force field selection and ligand parameterization. Moreover, a wider application of FEP methods has been seriously limited because of their very high computational demands, even for small size libraries. Recent breakthroughs in Machine Learning (ML) algorithms and big data mining along with an exponential growth of computing power, have led to promising applications of ML techniques in development of novel scoring functions.^10^ ML-based scoring functions have demonstrated remarkable performance in various benchmarking studies,^11^ and are in general several orders of magnitude faster than traditional scoring functions.

Regardless of classification, all scoring functions developed to date share a common obstacle that is they need a clear picture of explicit protein-ligand interactions (*i*.*e*., hydrogen bonds, polar and hydrophobic interactions, Van der Waals contacts, *etc*.). It implies a prerequisite slow and expensive sampling attempt prior to scoring calculation. Hence, even though modern ML-based scoring functions are significantly faster and more accurate, their implementation in a molecular docking pipeline could barely resolve the aforementioned drawbacks because of the vital sampling stage bottleneck.

In this study, we introduce a novel ML-based scoring function (iScore) that quickly and precisely predicts the binding affinity of protein-ligand complexes without the need for knowledge of explicit intermolecular interactions. Instead, iScore predicts the protein-ligand binding affinity based on a combination set composed of the ligand and binding pocket descriptors. Therefore, the sampling stage can be skipped, which leads to a massive saving in time and resources. On the other hand, since the iScore architecture is independent of explicit intermolecular interactions, it can be employed to score and rank a huge library of *de novo* small molecules against a protein target of interest, that greatly assists researchers to evaluate “unseen” regions of chemical space. iScore has been trained on the PDBbind 2020^12^ refined set using three different ML approaches: Deep Neural Network (*iScore-DNN*), Random Forest (*iScore-RF*), and eXtreme Gradient boosting (*iScore-XGB*). Furthermore, a hybrid scoring function (*iScore-Hybrid*) has been developed by combining and taking advantage of these three base-learners. The scoring power, ranking power, and screening power performances of iScore have been extensively tested and compared to other traditional and ML-based scoring functions using three different test sets: PDBbind 2016 core set (Comparative Assessment of Scoring Functions, CASF-2016),^4^ and two datasets from Community Structure-Activity Resource (CSAR NRC-HiQ Set1 and CSAR NRC-HiQ Set2).^13^ The authors believe that iScore opens the door to a new era of *de novo* drug discovery and pharmaceutics.

## 2. Materials and Methods

### 2.1. Dataset preparation

The iScore models have been trained on the PDBbind 2020 refined set as training dataset.^12^ The PDBbind 2020 refined set consists of 5,316 protein-ligand complexes along with the associated experimental affinity data and is a cherry-picked subset of the PDBbind 2020 general set with over 23,496 complexes, by selecting the complexes without any obvious structural issues or steric clashes, crystal resolution < 2.5 Å, R-factor < 0.25, non-covalent ligand binding, and affinity data reported as either *K*_*d*_ or *K*_*i*_ in the range of 10 mM to 1 pM. A full description of the criteria used for selecting the PDBbind refined set can be found in the original paper.^4^ The PDBbind 2016 core set, the first test set in our study and in the CASF-2016 benchmarking, was selected from the PDBbind refined set by applying even stricter criteria as follows: (1) The PDBbind refined set was subjected to a sequence similarity clustering with a similarity cutoff of 90% and only the clusters containing more than 5 members were considered, (2) five representative complexes were selected for each remaining cluster based on their affinity data, with the highest and lowest affinities having at least 100-fold difference, and three additional complexes, (3) the ligands should not be identical or stereoisomers throughout the PDBbind core set, (4) the electron density map and the ligand binding pose in each complex should be of high quality. It resulted in 285 protein-ligand complexes clustered into 57 clusters in the PDBbind 2016 core set. Other two test datasets used in this study are CSAR NRC-HiQ Set1 and CSAR NRC-HiQ Set2 containing 176, and 167 high quality protein-ligand complexes, respectively.

Prior to database preparation, the overlapping complexes between the PDBbind 2020 refined set and PDBbind 2016 core set were removed from the training set. In addition, the overlapping complexes between PDBbind 2020 refined set and CSAR NRC-HiQ Set1/Set2 were removed from the latter. The crystal structures were subsequently prepared using the PrepWizard in the Schrödinger 2023-2 program package (*https://www.schrodinger.com/*). Hydrogen atoms were incorporated, and missing side chain atoms were added using Prime. After fixing the potential structural defects, water molecules were removed from the complexes and the protonation states of ionizable residues were determined at pH = 7.0 by using PROPKA.^14^ The correct protonation states of the ligand molecules were determined at pH = 7.0 using Epik.^15^ The prepared complexes were further refined using the OPLS4 force field^16^ in a restrained minimization procedure with an RMSD threshold of 0.3 Å for all heavy atoms. The complexes which failed during the preparation stage were discarded. The final prepared datasets contain 4898 (PDBbind 2020 refined set), 285 (PDBbind 2016 core set), 68 (CSAR NRC-HiQ Set1), and 75 (CSAR NRC-HiQ Set2) complexesrespectively. The PDB codes in each dataset are listed in Table S1.

### 2.2. Descriptor Calculations

#### 2.2.1. Ligand descriptors

The 3D structures of the ligand molecules were converted to the corresponding canonical Simplified Molecular-Input Line-Entry System (SMILES)^17^ strings and subsequently a series of 81 1D/2D molecular descriptors were calculated using the RDKit library (https://www.rdkit.org) in Python such as: logarithm of partition coefficient (*MolLogP*), molecular refractivity (*MolMR*), exact molecular weight (*ExactMolWt*), number of heavy atoms (*HeavyAtomCount*), number of hydrogen bond acceptors (*NumHAcceptors*), number of hydrogen bond donors (*NumHDonors*), number of rotatable bonds (*NumRotatableBonds*), *etc*. A full list of the molecular descriptors used in this study is presented in Table S2. Figure S1 shows the histogram distribution of some molecular descriptors of the ligands in the training set along with the logarithmic form of the experimental binding affinity values (*pK*_*aff*_).

#### 2.2.2.. Binding pocket descriptors

The FPocket^18^ tool was employed to calculate 41 descriptors of the protein binding pocket such as pocket volume (*pock_vol*), number of alpha spheres (*nb_AS*), mean alpha sphere radius (*mean_as_ray*), mean alpha sphere solvent accessibility (*mean_as_solv_acc*), Polarity Score (*polarity_score*), Hydrophobicity Score (*hydrophobicity_score*), Charge Score (*charge_score*), Volume Score (*volume_score*), amino acid composition, *etc*. The protein binding pocket was explicitly defined by all atoms situated at a certain cutoff distance from the ligand molecule (3-7 Å). The initial assessments showed that a cutoff distance of 5 Å resulted in the best training and binding affinity prediction performance. A full list of the binding pocket descriptors is presented in Table S2. Figure S2 shows the histogram distribution of some descriptors of the binding pocket in the training dataset. Furthermore, FPocket suggests an intuitive estimation of the volume of potential ligands (*LigVol*_*BP*_) which was used in this study as a descriptor in training of the scoring models and as a key feature in training of the Ultra-Fast Screening (*UFS*) model which was used to improve the screening power performance by further filtering false positives (section 2.4.3).

### 2.3. Machine Learning Algorithms

iScore has been trained using three different ML approaches: Deep Neural Network (*DNN*),^19^ Random Forest (*RF*),^20^ and eXtreme Gradient boosting (*XGB*).^21^ In this study, the hyperparameters of the iScore-RF and iScore-XGB models were automatically tuned with the Bayesian optimization (BO)^22^ technique implemented in the Scikit-learn^23^ version 1.0.2, while Keras version 2.4.3 (https://keras.io)^24^ was employed for hyperparameter optimization of the iScore-DNN model. A 3-fold cross-validation was used to evaluate various hyperparameter combinations, and root mean squared error (RMSE) was utilized as the object function. The maximum number of iterations was set to 200.

#### 2.3.1. Deep Neural Network (DNN)

The Keras package version 2.4.3 in Python 3 was employed to build the iScore-DNN model. The DNN model consists of five layers: an input layer with 350 neural nodes, three hidden layers with 250, 150, and 50 neural nodes, and an output single node layer. The RELU^25^ activation function was used for all layers except the output layer where a LINEAR activation function was employed. The loss function and evaluation metric were set to Mean-Absolute-Error and Mean-Squared-Error, respectively. An Inverse-Time-Decay scheduler with an initial learning rate of 0.001, decay rate of 0.3, and decay steps of 8000 was used to properly lower the learning rate during the training process with Adam optimizer and 100 epochs. iScore-DNN was trained through 10×10-fold cross validation (XV) with random data shuffling in each XV loop. The final output was an average value over 100 DNN XV models.

#### 2.3.2. Random Forest (FR)

The Scikit-learn package version 1.0.2 in Python 3 was used to build the iScore-RF model. The random forest consisted of 200 decision trees (*n_estimators*) with *min_samples_split* (minimum number of samples required to split an internal node) = 2, *min_samples_leaf* (minimum number of samples required to be at a leaf node) = 1, *max_features* (number of features to consider when looking for the best split) = “auto”. The criterion was set to Mean-Squared-Error and the estimators were allowed to expand until all leaves were pure. iScore-RF was trained through 10×10-fold XV with a random data shuffling in each XV loop. The final output was an average value over 100 RF XV models.

#### 2.3.3. eXtreme Gradient Boosting (XGB)

The XGBoost package version 1.5.2 in Python 3 was used to build the iScore-XGB model. The hyperparameters of the XBG trainer are *n_estimators* (number of estimators) = 1000, *learning_rate* = 0.01, *subsample* (Subsample ratio of the training instances prior to growing estimators) = 0.7, *colsample_bytree* (subsample ratio of columns when constructing each tree)= 1.0, *max_depth* (maximum depth of an estimator) = 8, and *objective* (regression type) = “reg:squarederror”. iScore-XGB was trained through 10×10-fold XV with a random data shuffling in each XV loop. The final output was an average value over 100 XGB XV models.

#### 2.3.4. Hybrid Model

A hybrid scoring function (iScore-Hybrid) was developed by combining the iScore-DNN, iScore-RF, and iScore-XGB models. For this purpose, the average predicted affinity values over 100 XV of each model (iScore-DNN, iScore-RF, and iScore-XGB) along with experimental affinity data were fed into a DNN trainer. The iScore-Hybrid model consists of four layers: an input layer with 100 neural nodes, two hidden layers with 50 and 10 neural nodes, and an output single node layer. The RELU activation function was used for all layers except the output layer where a LINEAR activation function was employed. The loss function and evaluation metric were set to Mean-Absolute-Error and Mean-Squared-Error, respectively. An Inverse-Time-Decay scheduler with an initial learning rate of 0.001, decay rate of 0.3, and decay steps of 8000 was used to properly lower the learning rate during the training process with Adam optimizer and 100 epochs. iScore-Hybrid was trained through 10×10-fold XV with a random data shuffling in each XV loop. The final output was an average value over 100 DNN XV models.

### 2.4. Evaluation Metrics

#### 2.4.1. Scoring power

“S*coring power*” indicates the degree of correlation in the predicted versus experimentally determined binding affinity values. Hence, the Pearson correlation coefficient (R) was computed as a quantitative indicator of the scoring power (Eq. 1).^4^ The root mean squared error (RMSE) of the regression was also considered as additional indicator (Eq. 2).^4^

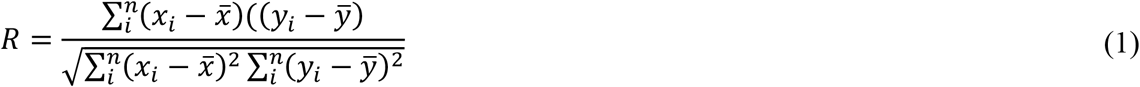

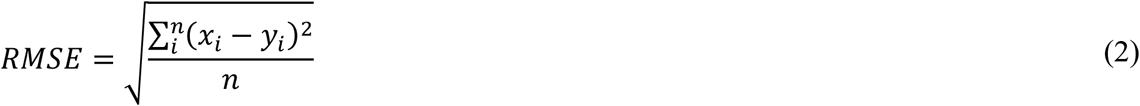

where, 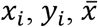 and 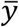 are the estimated and experimental binding affinities of the i^th^ complex and the corresponding average values, respectively. The summation upper limit (*n*) is the total number of complexes *i*.*e*., 4898 (PDBbind 2020 refined set), 285 (PDBbind 2016 core set), 68 (CSAR NRC-HiQ Set1), and 75 (CSAR NRC-HiQ Set2).

#### 2.4.2. Ranking power

“*Ranking power*” refers to the capability of the scoring function in ranking a given set of active ligands, with respect to their predicted binding affinity values, towards a particular protein target. The PDBbind 2016 core set contains 285 protein-ligand complexes clustered into 57 clusters. Each cluster contains a particular target receptor and 5 different active binders where the difference between binding affinities of the strongest and weakest binders is at least 100-fold. Figure S3 shows a boxplot of experimental binding affinity values for each of the 57 clusters in the PDBbind 2016 core set. The Spearman ranking correlation (*ρ*, Eq. 3)^4^ was used as an indicator of the ranking power (as in the CASF-2016 benchmarking) since in contrast to scoring power, ranking power does not request a linear correlation between experimental and predicted binding affinity values.

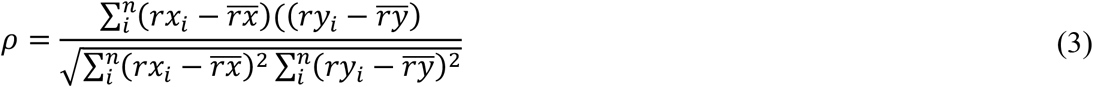

where, 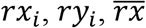 and 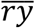 are the rank of the estimated and experimental binding affinities of the i^th^ complex and the corresponding average values, respectively. The summation upper limit (*n*) is the total number of samples in each cluster, that is five in this case. The average Spearman ranking correlation, < *ρ* >, was subsequently calculated over all 57 target proteins in the PDBbind 2016 core set.

#### 2.4.3. Screening power

“*Screening power*” indicates the ability of a scoring function to identify the true binder with the highest affinity against a given protein target among a set of random decoy molecules. The first quantitative reference metric of screening power is the success rate of identifying the ligand with highest affinity against each 57 target receptors, in the PDBbind 2016 core set, among the 1, 5, and 10% predicted top candidates. The second indicator is the success rate of identifying all binders with experimental binding affinity values less than 10 mM (*pK*_*d*_ ≥ 2), 10 µM (*pK*_*d*_ ≥ 5), 1 µM (*pK*_*d*_ ≥ 6), 0.1 µM (*pK*_*d*_ ≥ 7), 0.01 µM (*pK*_*d*_ ≥ 8), and 1 nM (*pK*_*d*_ ≥ 9), among the 1, 2, 3, 5, and 10% top candidates over all 285 complexes. There are 285, 213, 167, 117, 75, and 39 binders with experimental binding affinity values less than 10 mM, 10 µM, 1 µM, 0.1 µM, 0.01 µM, and 1 nM in the PDBbind 2016 core set, respectively.

The screening power performance of the iScore models was further improved by a “Ultra-Fast Screening” (*UFS*) stage prior to the binding affinity prediction. In the *UFS* stage, the ligands that volumetrically do not match with a given receptor’s binding pocket (too big or too small) will be filtered out. One of the features that FPocket tool predicts, after receptor’s binding pocket evaluation, is an intuitive estimation of the volume of potential binders (*LigVol*_*BP*_) that strongly correlates with the receptor’s binding pocket volume (*pock_vol*) (Figure 1a). Hence, an RF-based regression model was trained to calculate *LigVol*_*BP*_ based on 2D molecular descriptors (*LigVol*_*pred*_). From the correlation graph, 99% prediction band (Figure 1b) was calculated upon 3×10-fold XV and was used in the UFS stage, so that only the ligands with predicted volumes (*LigVol*_*pred*_) within the 99% prediction band of the *LigVol*_*BP*_ value were allowed to pass to the scoring stage. The trained RF volume-predictor was tested on three test sets used in this study (PDBbind 2016 core set, CSAR NRC-HiQ Set1 and Set2) and the results show a very close to perfect correlations (Figures 1c – 1e).

**Figure 1.**
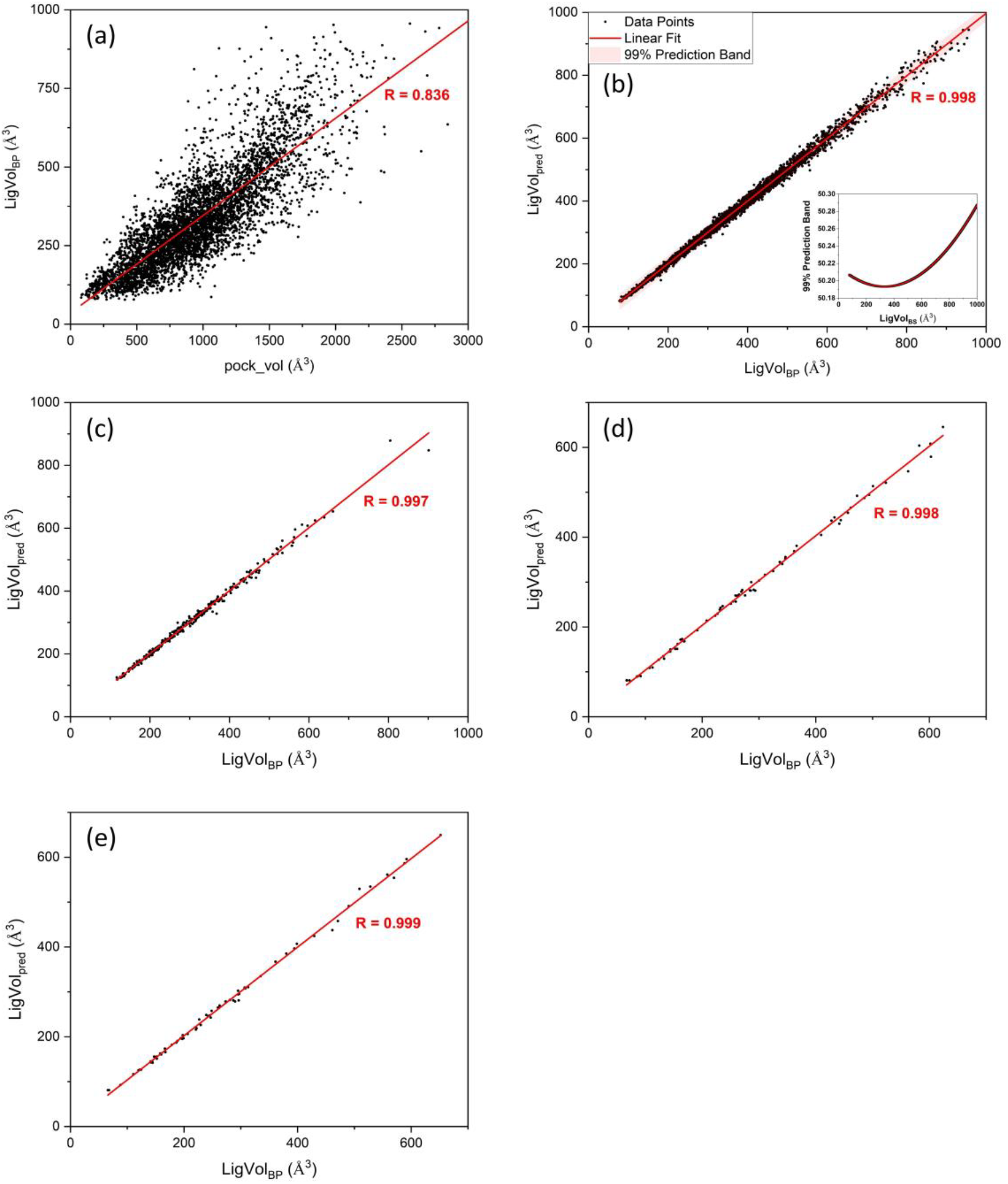
**(a)** The strong correlation between the *LigVol*_*BP*_ and *pock_vol*. **(b)** The correlation between *LigVol*_*pred*_ estimated from 2D molecular descriptors and *LigVol*_*BP*_ with 99% prediction band region calculated upon 3×10-fold XV on the training set. The performance of the RF volume predictor on **(c)** the PDBbind 2016 core set, **(d)** the CSAR NRC-HiQ Set1 and **(e)** the CSAR NRC-HiQ Set1. The Pearson correlation coefficients (*R*) are shown in each graph.

### 2.5. Results

#### 2.5.1. Training

Figures 2a-2d show the 25%-75% boxplot presentation and distribution of Pearson (*R*) and Spearman (*ρ*) correlation coefficients along with the mean, median and standard deviation (SD) values for the models trained with different ML algorithms (base-learners) and the hybrid model, upon 10×10-fold XV training campaign. The mean Pearson coefficients are 0.75, 0.75, 0.77, and 0.78 for the iScore-DNN, iScore-RF, iScore-XGB, and iScore-Hybrid models respectively. The SD profile of the Pearson coefficient is similar for all models and lies in the range of 0.01– 0.02. As the figures indicate, the mean Spearman coefficients (*ρ*) are slightly lower than the mean Pearson coefficients, but the same SD values have been observed. Figure 2e shows the RMSE statistics for three base-learners along with the hybrid model. The mean RMSE values are 1.32, 1.30, 1.25, and 1.23 for iScore-DNN, iScore-RF, iScore-XGB, and iScore-Hybrid models, respectively. The SD profiles of the RMSE metric are similar and cover the range of 0.04 – 0.05. The results clearly show that iScore-Hybrid outperforms the base-learners with higher mean Pearson and Spearman correlation coefficients and lower mean RMSE value upon the cross-validation training campaign.

**Figure 2.**
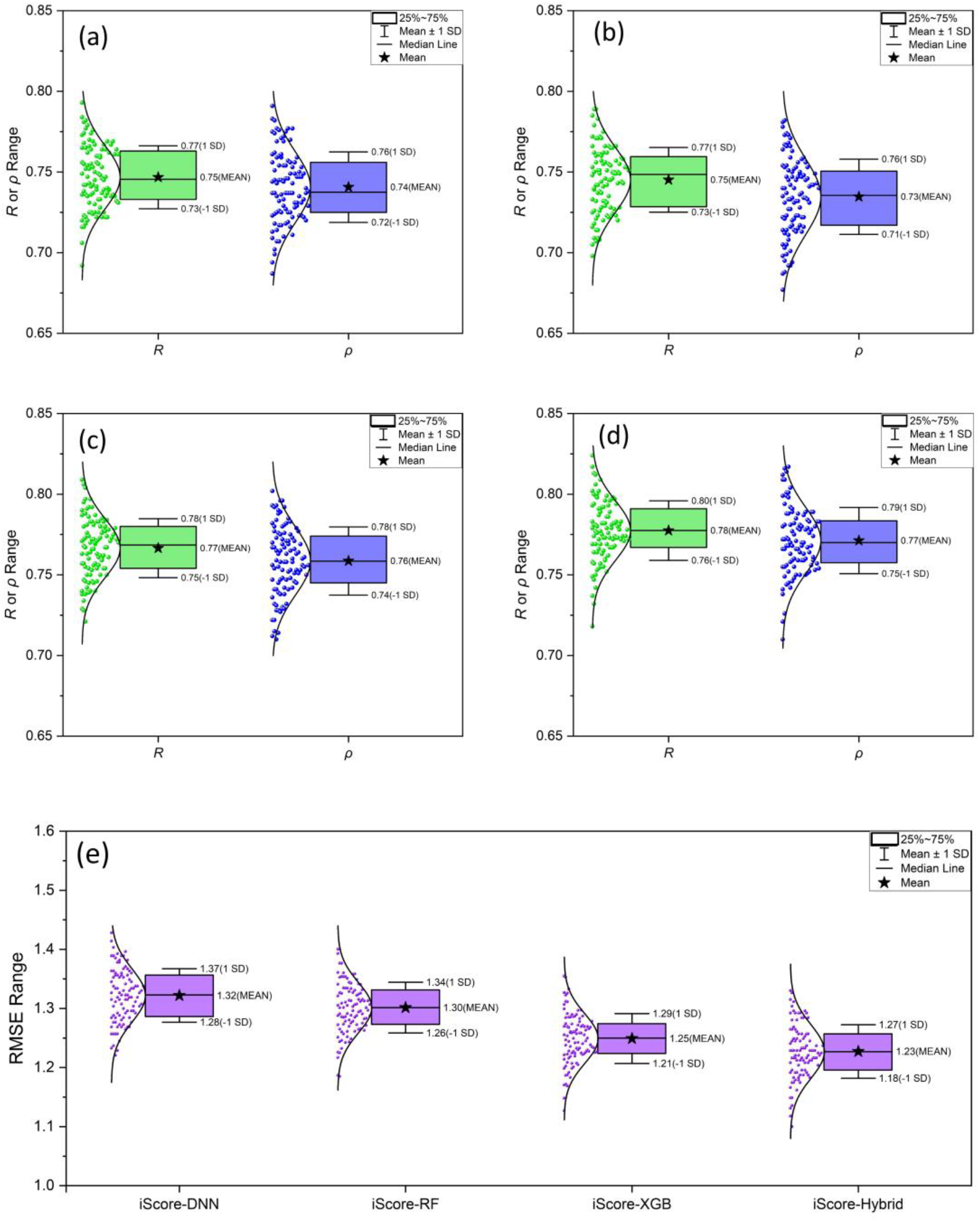
Boxplot presentation and distribution of Pearson (*R;* green) and Spearman (*ρ*; blue) correlation correlations along with the mean, median and standard deviation (SD) values for the **(a)** iScore-DNN, **(b)** iScore-RF, **(c)** iScore-XGB, and **(d)** iScore-Hybrid upon 10×10-fold XV training campaign. **(e)** The RMSE statistics for the base-learners along with the hybrid model.

To further understand the better performance of iScore-Hybrid over the three base-learners, the plots of the *squared error* (squared difference between the experimental and the predicted *pK*_*aff*_) versus the experimental *pK*_*aff*_ (Figure 3) associated to each model, have been deeply explored. As Figure 3 illustrates, there are three distinct regions which are quantitatively distinguished after fitting the data into a Piecewise Linear function with three segments (PWL3). The first region (green area) is the trust-zone in the mid-range *pK*_*aff*_ spectrum where the PWL3 function forms a horizontal line indicating the most reliable range of *pK*_*aff*_ that the model can predict at the maximum accuracy (minimum error). The other two regions are at the two ends of the experimental *pK*_*aff*_ (yellow areas) where the PWL3 function forms nonzero-slope lines. One can elucidate the overall performance of the models by comparing three determinative factors: the trust-zone’s length (the bigger the better) and height (the lower the better) and the absolute slope of the lines in the nonzero-slope regions (the lower the better). The maximum trust-zone’s length is 5.82 [3.37, 9.19] and is found for iScore-DNN, while the corresponding value is 3.30 [4.64, 7.94], 3.45 [4.55, 8.00], and 3.91 [4.20, 8.11] for iScore-RF, iScore-XGB, and iScore-Hybrid, respectively. One the other hand, iScore-DNN has the maximum trust-zone’s height of 1.53 followed by 1.12 (iScore-Hybrid), 0.99 (iScore-XGB), and 0.97 (iScore-RF). Moreover, the minimum absolute slope of the lines in the nonzero-slope regions belong to iScore-Hybrid which are 1.60 and 0.98 at the low and high affinity limits, respectively. The corresponding values are (1.63 and 1.38), (2.00 and 1.54), and (3.27 and 2.05), for iScore-XGB, iScore-RF, and iScore-DNN, respectively. Therefore, in the context of the trust-zone length, iScore-Hybrid showed a better performance than two base-learners iScore-RF and iScore-GXB. In the context of the trust-zone height, iScore-Hybrid outperforms iScore-DNN by a considerable margin. Interestingly, iScore-Hybrid also showed the best performance at the low and high affinity limits where the base-leaners suffer from lower prediction accuracy.

**Figure 3.**
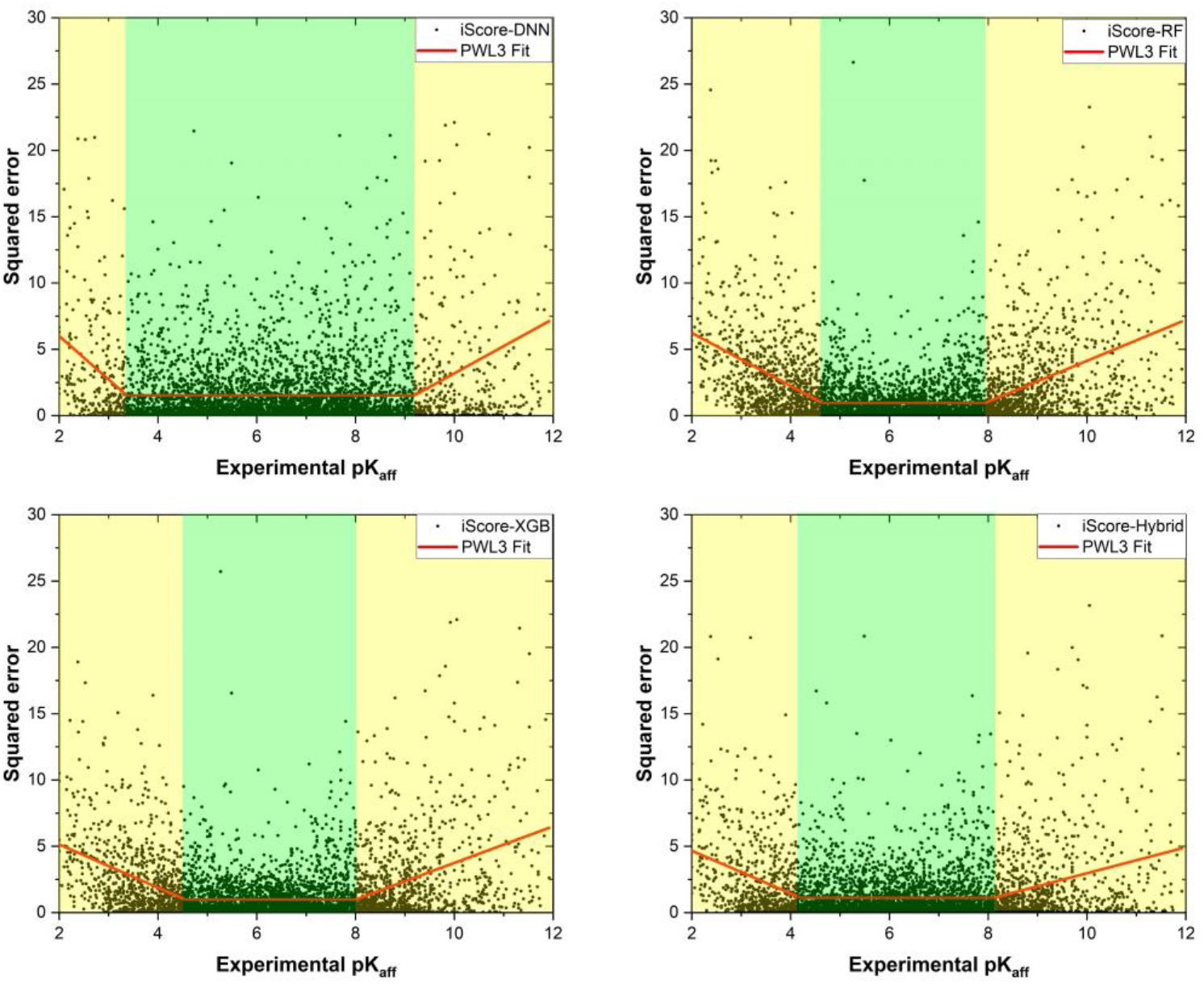
S*quared error* versus experimental *pK*_*aff*_ obtained from each model upon 10×10-fold XV training. The Piecewise Linear function with three segments (PWL3) fitted into the data are shown in red. The green and yellow areas illustrate the trust-zone and nonzero-slope regions, respectively.

#### 2.5.2. Benchmarks

The scoring, ranking, and screening power performances of the iScore models were extensively tested and compared to other traditional and ML-based scoring functions on three different test sets: PDBbind 2016 core set (scoring, ranking, and screening power), and CSAR NRC-HiQ Set1 and Set2 (scoring performance).

##### 2.5.2.1. Scoring power

Figures 4a and 4b illustrate the scoring power performance of the iScore models versus 40 traditional and modern ML-based scoring functions in terms of the Pearson correlation coefficient and RMSE metrics tested on the PDBbind 2016 core set, respectively. As these figures show, iScore-Hybrid outperforms the base-learners in the context of both scoring power metrics (*R* = 0.814 and *RMSE* = 1.34) and stands among the top scoring functions. Two major competitors are *graphDelta*^26^ and *K*_*deep*_^27^. *graphDelta* is a ML graph-based scoring function that employs a message passing neural network (MPNN) for modeling protein−ligand interactions. *graphDelta* yielded the best scoring power metrics on the PDBbind 2016 core set with *R* = 0.87 and *RMSE* = 1.05. *K*_*deep*_ is also an ML-based scoring function which uses a 3D-convolutional neural network for predicting the ligand binding affinities. *K*_*deep*_ demonstrated a very good scoring power performance on the PDBbind 2016 core set with *R* = 0.82 and *RMSE* = 1.27. Nonetheless, these scoring functions, like any other scoring functions published up to now, require a full picture of the protein-ligand interactions which imposes critical limitations on their speed and applicability as discussed earlier. Figures 4c and 4d compare the Pearson correlation coefficient and *RMSE* metrics of the iScore models against modern ML-based scoring functions, tested on the CSAR NRC-HiQ Set1and Set2, respectively. While iScore-Hybrid is amongst the top 3 best performing scoring functions on the PDBbind 2016 core set, it is at the top of the list when tested on the CSAR NRC-HiQ Set1 (*R* = 0.834 and *RMSE* = 1.27) and Set2 (*R* = 0.767 and *RMSE* = 1.32). The iScore base-learners also outperform the other scoring functions including *graphDelta* (*R* = 0.74, 0.65 and *RMSE* = 1.59, 1.52) and *K*_*deep*_ (*R* = 0.72, 0.75 and *RMSE* = 2.08, 1.91). As the test results indicate (Figure 4), iScore-XGB and iScore-DNN are the best performing base-learners in terms of the Pearson correlation coefficient and *RMSE* metrics on all three test sets, respectively.

**Figure 4.**
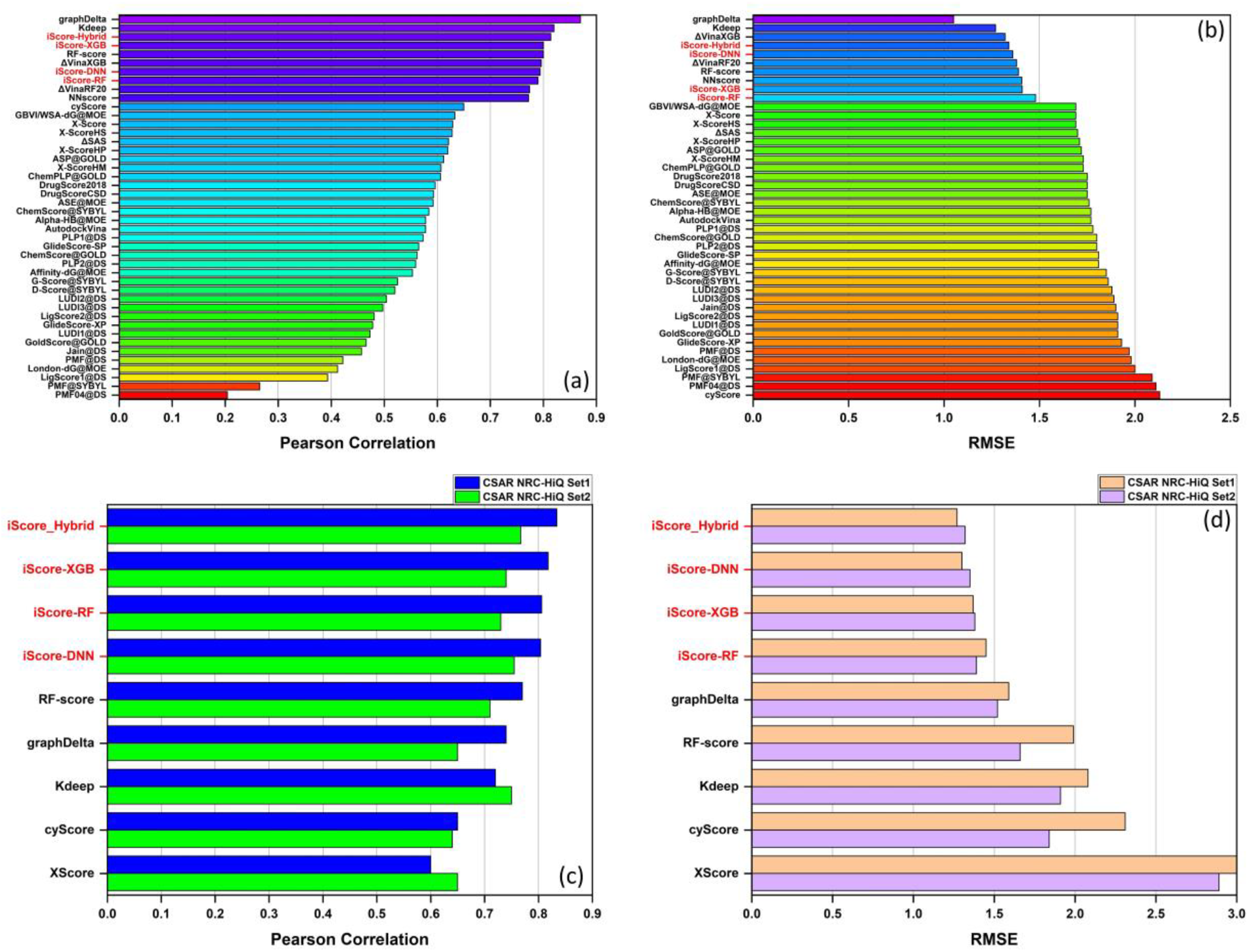
Scoring power performance of the iScore models compared to other scoring functions tested on (a, b) the PDBbind 2016 core set; and (c, d) the CSAR NRC-HiQ Set1 and Set2. Scoring functions are ranked by the Pearson correlation coefficients in descending order.

Figure 5 illustrates the *squared error* (squared difference between the experimental and the predicted *pK*_*aff*_) versus the experimental *pK*_*aff*_ associated to each iScore model obtained on the PDBbind 2016 core set. The trust-zone’s lengths of iScore-Hybrid and iScore-DNN are similar (∼ 2.1 [6.0, 8.1]) while iScore-RF and iScore-XGB showed a higher value (∼ 2.5 [5.5, 8.0]). One the other hand, iScore-Hybrid has the minimum trust-zone’s height of 0.48 followed by 0.56 (iScore-DNN), 0.59 (iScore-RF), and 0.60 (iScore-XGB). Furthermore, iScore-Hybrid shows the best performance at the low and high affinity limits where the absolute slope of the lines in these regions are 1.10 and 1.67, respectively. The corresponding values are (1.12 and 1.72), (1.68 and 2.48), and (1.56 and 2.06), for iScore-DNN, iScore-RF, and iScore-XGB, respectively. Therefore, except the trust-zone’s length, iScore-Hybrid outperforms the base-learners.

**Figure 5.**
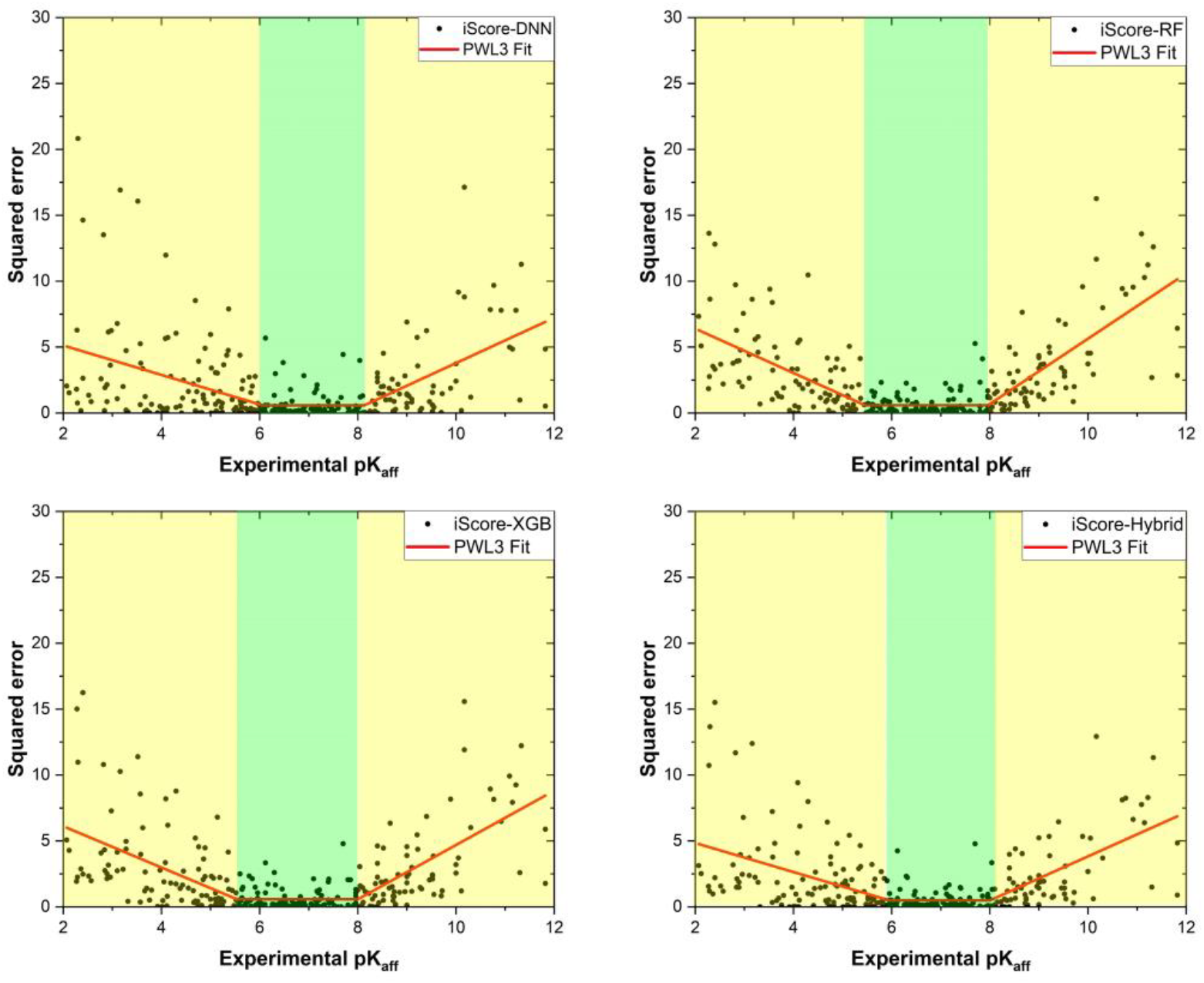
The *squared error* versus the experimental *pK*_*aff*_ obtained from each model upon testing on the PDBbind 2016 core set. The Piecewise Linear function with three segments (PWL3) fitted into the data are shown in red. The green and yellow areas illustrate the trust-zone and nonzero-slope regions, respectively.

##### 2.5.2.2. Ranking power

Figures 6a-6d show the disaggregated and average (vertical red dashed lines) ranking Spearman correlation coefficients ( *ρ* ) over the 57 targets in the PDBbind 2016 core set evaluated by the iScore base-learners and the hybrid model. Figure 6e illustrates the raking power performance of the iScore models (based on the average Spearman correlation coefficient) and compares those with several other scoring functions. As this figure shows, iScore-Hybrid (<*ρ*> = 0.705) outperforms not only the base-learners but all other scoring functions in the ranking campaign. As Figure 6e indicates, iScore-Hybrid is followed by iScore-RF (<*ρ*> = 0.702), iScore-DNN (<*ρ*> = 0.691), and iScore-XGB (<*ρ*> = 0.690), respectively. It is worth mentioning that the ranking power performances of iScore models are significantly better than *K*_*deep*_ (<*ρ*> = 0.51) which was one of the major competitors in the scoring power campaign.

**Figure 6.**
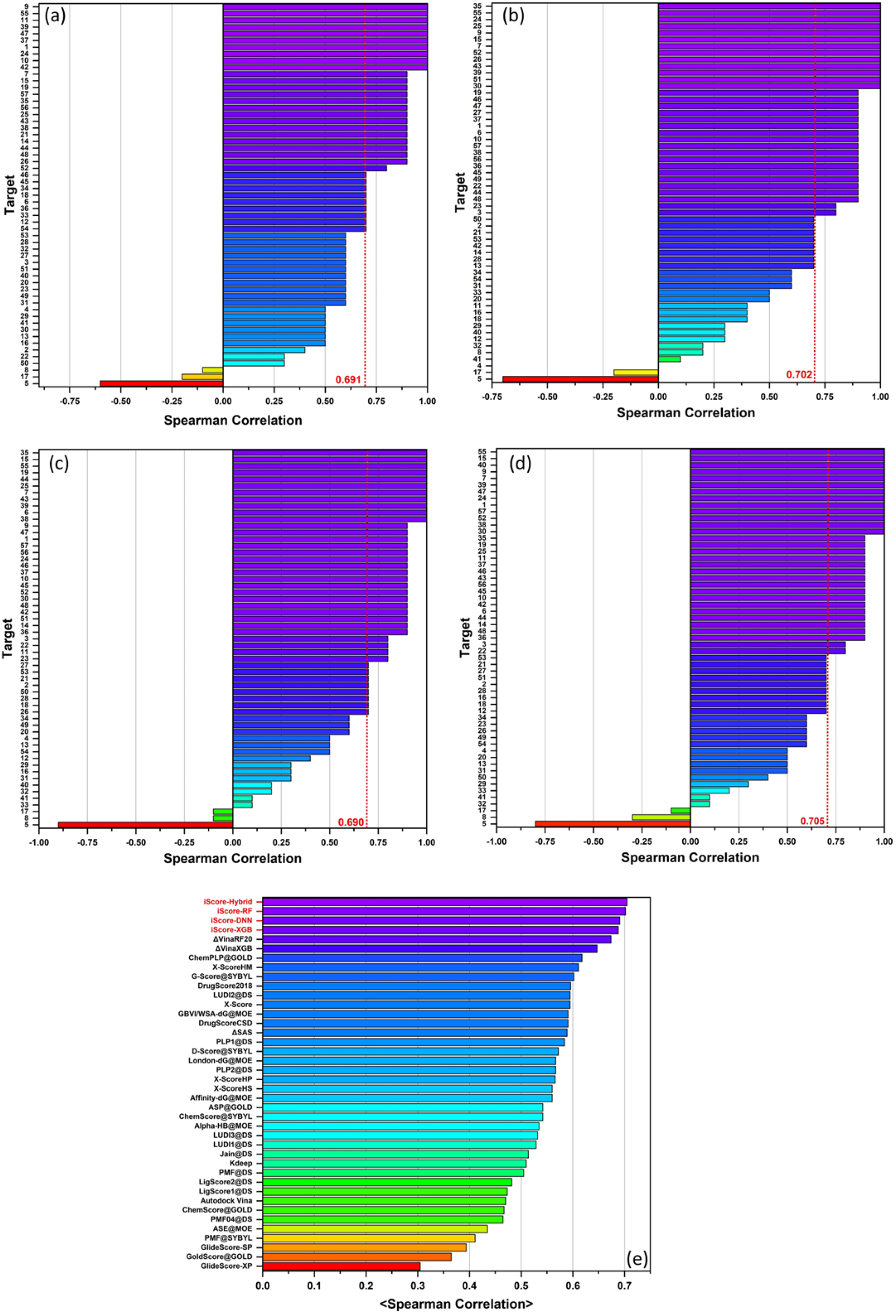
The ranking power performance of (a) iScore-DNN, (b) iScore-RF, (c)-iScore-XGB, and (d) iScore-Hybrid on the CASF-2016 test set based on the Spearman correlation coefficient of individual targets. (e) A comparison between the ranking power performance of the iScore models and other scoring functions based on the average Spearman correlation coefficients (vertical red dashed lines in (a)-(d)) over all targets.

##### 2.5.2.3. Screening power

Figure 7(a) shows the screening power performance of iScore in terms of success rate of identifying the highest affinity ligand of each of the 57 target receptors in the PDBbind 2016 core set, among the 1, 5, and 10% top candidates. Figure 7(a) demonstrates that the screening performance of iScore (73.7% for iScore-Hybrid, iScore-XGB, and iScore-NDD and 68.4% for iScore-RF) is considerably better than all other scoring functions in the screen power campaign. Figure 7(b) illustrates the success rate of identifying all binders with the experimental binding affinity values less than 10 mM (*pK*_*d*_ ≥ 2), 10 µM (*pK*_*d*_ ≥ 5), 1 µM (*pK*_*d*_ ≥ 6), 0.1 µM (*pK*_*d*_ ≥ 7), 0.01 µM (*pK*_*d*_ ≥ 8), and 1 nM (*pK*_*d*_ ≥ 9), among the 1, 2, 3, 5, and 10% top candidates over all 285 complexes in the PDBbind core set. As this figure shows, the success rate of iScore increases as the *pK*_*d*_ of the binder increases. For instance, the success rate of iScore-Hybrid is 51.9, 61.5, 68.9, 81.2, 85.3, and 92.3% for identifying all binders with experimental binding affinity values less than 10 mM, 10 µM, 1 µM, 0.1 µM, 0.01 µM, and 1 nM, among the 10% top candidates.

**Figure 7.**
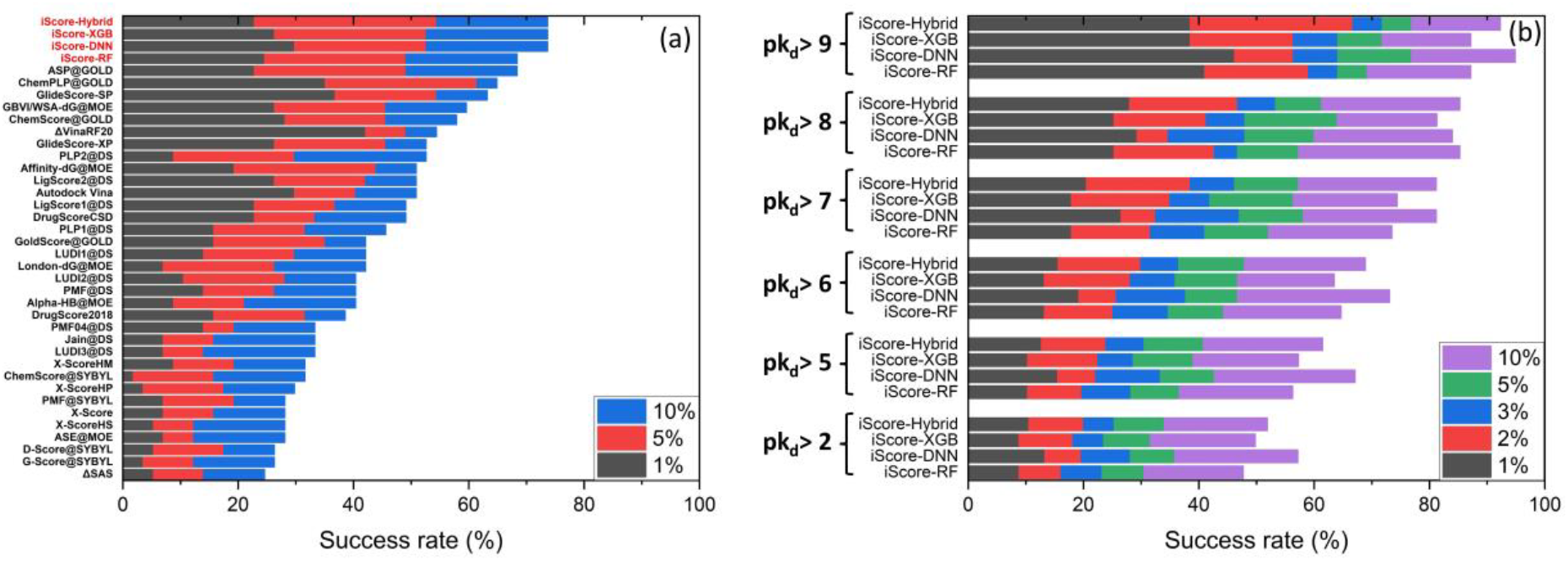
(a) Comparison between screening power performance of iScore and other scoring functions in success rate of identifying the highest affinity ligand of each of the 57 target receptors in the PDBbind 2016 core set, among the 1, 5, and 10% top candidates. (b) the success rate of identifying all binders with experimental binding affinity values less than 10 mM (*pK*_*d*_ ≥ 2), 10 µM (*pK*_*d*_ ≥ 5), 1 µM (*pK*_*d*_ ≥ 6), 0.1 µM (*pK*_*d*_ ≥ 7), 0.01 µM (*pK*_*d*_ ≥ 8), and 1 nM (*pK*_*d*_ ≥ 9), among the 1, 2, 3, 5, and 10% top candidates over all 285 complexes in the PDBbind core set.

#### 2.5.3. Speed performance

iScore was trained using a single compute node on the Alvis supercomputer allocated by the C3SE supercomputing facility with one NVIDIA Tesla A100 HGX GPU (40GB RAM), 32 core Intel(R) Xeon(R) Gold 6338 CPU @ 2GHz, and 256GB DDR4 RAM. iScore is capable of screening >8000 compound/s (∼700 million screenings a day) on a single compute node: 32 core AMD Ryzen 9 7950X CPU, NVIDIA RTX A4000 GPU (16GB RAM), and 32GB DDR5 RAM.

### 2.6. Conclusions

This work introduces iScore, a cutting-edge ML-based scoring function designed to predict the binding affinity of protein-ligand complexes with unprecedented precision and speed. Unlike traditional scoring functions that rely heavily on the explicit knowledge of intermolecular interactions, iScore leverages a novel approach. It utilizes a combination of ligand and binding pocket descriptors, thereby bypassing the need for extensive conformational sampling. This methodological innovation not only saves significant computational time and resources but also provides the applicability to evaluate vast molecular libraries, offering a leap towards a more efficient exploration of the chemical space. The benchmarking of iScore across multiple datasets highlights its robustness and superior performance over traditional and advanced scoring functions. Notably, the development of the hybrid iScore model (iScore-Hybrid), which integrates the strengths of individual base learners, sets new benchmarks in scoring, ranking, and screening capabilities essential for drug discovery processes. The innovation of iScore is further underscored by its practical implications. The ability to screen over 8000 compounds per second on a single GPU translates to the screening of 700 million compounds daily, illustrating the scalability and efficiency of iScore in handling ultra-large molecular libraries. This capability is critical in accelerating the drug discovery process, from initial screening to identification of lead compounds. The promising results of iScore not only sets a new standard in scoring function development but also open a new era in the utilization of machine learning technologies for drug discovery.

## Data Availability

The training datasets, pre-trained models, and instruction for the retraining is available by contacting the authors.

## Supporting Information

Additional information regarding the pdb codes used for the training and benchmarking, database analysis, and molecular and binding pocket descriptors are available in the supporting information.

## Acknowledgments

This research was funded by the EU’s Horizon 2020 research and innovation program under the MSCA-RISE program 734749 (INSPIRED) (L.A. Eriksson). The Faculty of Science at the University of Gothenburg and the Swedish Science Research Council (VR; grant no. 2019-3684), and the Swedish Cancer Foundation (CF; grant no. 21-1447-Pj) (L.A. Eriksson) are also gratefully acknowledged for funding. The authors thank the Swedish National Infrastructure for Computing/National Academic Infrastructure for Supercomputing in Sweden for generous allocations of computing time at supercomputing centers C3SE and NSC in part funded by the Swedish Research Council through grant agreement no. 2022-06725. The authors gratefully acknowledge André Stadelmann at ANYO Labs AB for professional parallelization of the tool to maximize the ultimate performance.

