## Supplementary Tables S1 and S2, and Figures S1, S2 and S3. for "iScore: A ML-Based Scoring Function for *de novo* Drug Discovery"


**Table S1.** PDB codes of the protein-ligand complexes used in each training and test database.

| Dataset | PDB codes |
|---|---|
| PDBbind 2020 refined set (Training set) | 1a1e, 1a4k, 1a4r, 1a4w, 1a9m, 1a9q, 1a28, 1a69, 1a94, 1a99, 1aaq, 1add, 1adl, 1ado, 1afk, 1afl, 1ai4, 1ai5, 1ai7, 1aid, 1aj7, 1ajn, 1ajp, 1ajq, 1ajv, 1ajx, 1alw, 1amk, 1amw, 1apv, 1atl, 1atr, 1avn, 1ax0, 1azm, 1b0h, 1b1h, 1b2h, 1b3f, 1b3g, 1b3h, 1b3l, 1b4h, 1b4z, 1b05, 1b5h, 1b5i, 1b5j, 1b6h, 1b7h, 1b8n, 1b8o, 1b8y, 1b9j, 1b32, 1b38, 1b40, 1b46, 1b51, 1b52, 1b55, 1b57, 1b58, 1bai, 1bdq, 1bgq, 1bhf, 1bhx, 1bju, 1bjv, 1bm7, 1bma, 1bn1, 1bn3, 1bn4, 1bnn, 1bnq, 1bnt, 1bnu, 1bnv, 1bnw, 1bp0, 1bq4, 1br6, 1bty, 1bv7, 1bv9, 1bwa, 1bwb, 1bxo, 1bxq, 1bxr, 1bzj, 1bzy, 1c1r, 1c1u, 1c1v, 1c3x, 1c4u, 1c5c, 1c5n, 1c5o, 1c5p, 1c5q, 1c5s, 1c5t, 1c5x, 1c5y, 1c70, 1c83, 1c84, 1c86, 1c87, 1c88, 1cbx, 1ceb, 1cet, 1cgl, 1ciz, 1cnw, 1cnx, 1cny, 1cps, 1ctt, 1ctu, 1d2e, 1d3d, 1d3p, 1d4h, 1d4i, 1d4j, 1d4p, 1d4y, 1d6v, 1d6w, 1d7i, 1d7j, 1d09, 1d9i, 1det, 1df8, 1dgm, 1dhi, 1dhj, 1dif, 1dl7, 1dmp, 1dqn, 1drj, 1drk, 1drv, 1dud, 1duv, 1dy4, 1dzk, 1e1v, 1e1x, 1e2k, 1e2l, 1e3g, 1e3v, 1e4h, 1e5j, 1e6q, 1e6s, 1eb2, 1ebw, 1ebz, 1ec0, 1ec1, 1ec2, 1ec3, 1ec9, 1ecq, 1ecv, 1efy, 1egh, 1ejn, 1ela, 1elb, 1elc, 1eld, 1ele, 1elr, 1enu, 1eoc, 1epo, 1erb, 1ew8, 1ex8, 1ez9, 1ezq, 1f0r, 1f0s, 1f0t, 1f0u, 1f3e, 1f4e, 1f4f, 1f4g, 1f4x, 1f5k, 1f5l, 1f8b, 1f8c, 1f8d, 1f8e, 1f57, 1f73, 1f74, 1fao, 1fch, 1fcx, 1fcy, 1fcz, 1fd0, 1fh7, 1fh8, 1fh9, 1fhd, 1fiv, 1fjs, 1fkb, 1fkf, 1fkg, 1fkh, 1fki, 1fkn, 1fkw, 1fl3, 1fm9, 1fo0, 1fpc, 1fq5, 1ftm, 1fv0, 1fzj, 1fzk, 1fzm, 1fzo, 1fzq, 1g1d, 1g2l, 1g2o, 1g3d, 1g3e, 1g4o, 1g7f, 1g7g, 1g7q, 1g7v, 1g30, 1g32, 1g35, 1g36, 1g45, 1g46, 1g48, 1g52, 1g53, 1g54, 1g74, 1g85, 1g98, 1gaf, 1gai, 1gar, 1gfy, 1ghv, 1ghw, 1ghy, 1ghz, 1gi1, 1gi4, 1gi7, 1gj6, 1gjc, 1gnm, 1gnn, 1gno, 1grp, 1gvw, 1gvx, 1gwv, 1gx8, 1gyx, 1gyy, 1h0a, 1h1s, 1h2k, 1h2t, 1h4w, 1h5v, 1h6h, 1h46, 1hbv, 1hdq, 1hee, 1hfs, 1hi3, 1hi4, 1hi5, 1hih, 1hii, 1hk4, 1hlk, 1hmr, 1hms, 1hmt, 1hn4, 1hos, 1hp5, 1hpo, 1hps, 1hpv, 1hpx, 1hsh, 1hsl, 1hvh, 1hvi, 1hvj, 1hvk, 1hvl, 1hvr, 1hvs, 1hwr, 1hxb, 1hxw, 1hyo, 1i1e, 1i2s, 1i5r, 1i7z, 1i9n, 1i9p, 1i37, 1ie9, 1if7, 1if8, 1igb, 1igj, 1ii5, 1iih, 1iiq, 1ik4, 1ikt, 1ivp, 1iy7, 1izh, 1izi, 1j01, 1j4r, 1j14, 1j16, 1j17, 1j36, 1j37, 1jak, 1jao, 1jaq, 1jcx, 1jet, 1jeu, 1jev, 1jgl, 1jlr, 1jmf, 1jmg, 1jn4, 1jq8, 1jqy, 1jsv, 1jvu, 1jyq, 1jys, 1jzs, 1k1j, 1k1l, 1k1m, 1k1n, 1k1o, 1k1y, 1k4g, 1k4h, 1k6c, 1k6p, 1k6t, 1k6v, 1k9s, 1k21, 1k22, 1k27, 1kav, 1kc7, 1kdk, 1kel, 1kjr, 1km3, 1kmy, 1koj, 1kpm, 1ksn, 1kug, 1kui, 1kuk, 1kv1, 1kv5, 1kyv, 1kzk, 1kzn, 1l8g, 1l83, 1laf, 1lag, 1lah, 1lbf, 1lbk, 1lee, 1lf2, 1lgt, 1lgw, 1lhu, 1li2, 1li3, 1li6, 1lke, 1lkk, 1lkl, 1lnm, 1loq, 1lpk, 1lpz, 1lrh, 1lst, 1lvu, 1lyb, 1lyx, 1lzq, 1m0b, 1m0n, 1m0o, 1m0q, 1m1b, 1m2p, 1m2q, 1m2r, 1m2x, 1m4h, 1m5w, 1m7i, 1m48, 1m83, 1mai, 1mes, 1met, 1mfa, 1mfd, 1mfi, 1mjj, 1mmq, 1mmr, 1moq, 1mq5, 1mrn, 1mrs, 1mrw, 1mrx, 1msm, 1msn, 1mu6, 1mu8, 1mue, 1my4, 1n0s, 1n1m, 1n3i, 1n4h, 1n4k, 1n5r, 1n46, 1n51, 1ndv, 1ndw, 1ndy, 1ndz, 1nf8, 1nfu, 1nfw, 1nfx, 1nfy, 1nh0, 1nhu, 1nhz, 1nja, 1njc, 1njd, 1nje, 1njs, 1nl9, 1nli, 1nm6, 1nny, 1no6, 1np0, 1nq7, 1nt1, 1nvr, 1nvs, 1nw4, 1nw5, 1nw7, 1nwl, 1nz7, 1o0f, 1o0m, 1o0n, 1o1s, 1o2h, 1o2j, 1o2n, 1o2o, 1o2q, 1o2r, 1o2w, 1o2z, 1o3d, 1o3i, 1o3j, 1o3l, 1o5a, 1o5c, 1o5e, 1o5g, 1o5r, 1o7o, 1o30, 1o33, 1o35, 1o36, 1o38, 1o86, 1oar, 1oau, 1oba, 1ocq, 1od8, 1odi, 1odj, 1oe8, 1ogd, 1ogg, 1ogx, 1ogz, 1ohr, 1oif, 1okl, 1om1, 1ony, 1onz, 1ork, 1os0, 1os5, 1oss, 1owe, 1oxr, 1oyq, 1oz0, 1p1o, 1p19, 1p57, 1pa9, 1pb8, 1pb9, 1pbq, 1pdz, 1pfu, 1pgp, 1phw, 1pkx, 1pme, 1pot, 1ppc, 1pph, 1ppi, 1ppk, 1ppl, 1ppm, 1pr5, 1pro, 1pvn, 1px4, 1pxo, 1pxp, 1pyn, 1pz5, 1pzi, 1pzp, 1q1g, 1q7a, 1q8w, 1q54, 1q65, 1q72, 1q84, 1q91, 1qan, 1qaw, 1qb1, 1qb6, 1qb9, 1qbn, 1qbo, 1qbq, 1qbr, 1qbs, 1qbt, 1qbu, 1qbv, 1qf0, 1qf2, 1qft, 1qhc, 1qin, 1qji, 1qk4, 1qka, 1qkb, 1ql7, 1ql9, 1qxk, 1qxl, 1qy1, 1qy2, 1qyg, 1r0p, 1r1h, 1r1j, 1r9l, 1rbp, 1rd4, 1rjk, 1rmz, 1rnm, 1rnt, 1ro6, 1rp7, 1rpf, 1rpj, 1rr6, 1rtf, 1s5z, 1s19, 1s39, 1s89, 1sb1, 1sbg, 1sdt, 1sdu, 1sdv, 1sgu, 1sh9, 1siv, |

| | |
|---|---|
| | 1sl3, 1sld, 1sln, 1sqo, 1sqt, 1sr7, 1srg, 1ssq, 1stc, 1str, 1sv3, 1sw2, 1swg, 1swr, 1syh, 1szd, 1t4v, 1t5f, 1t7d, 1t7j, 1t31, 1t32, 1ta6, 1tcx, 1td7, 1thz, 1tjp, 1tkb, 1tlp, 1tmn, 1tng, 1tnh, 1tni, 1tom, 1tpw, 1tq4, 1trd, 1tsy, 1ttm, 1tx7, 1txr, 1u0g, 1u1w, 1u33, 1u71, 1ua4, 1ucn, 1ugx, 1uho, 1ui0, 1uj5, 1uml, 1uou, 1upf, 1ur9, 1usi, 1usk, 1usn, 1utj, 1utl, 1utm, 1utn, 1uv6, 1uw6, 1uwf, 1uwt, 1uwu, 1uz1, 1uz4, 1uz8, 1v0k, 1v0l, 1v1j, 1v2j, 1v2k, 1v2l, 1v2n, 1v2o, 1v2r, 1v2s, 1v2t, 1v2u, 1v2w, 1v7a, 1v48, 1vfn, 1vyf, 1vyg, 1vzq, 1w0z, 1w3j, 1w3k, 1w3l, 1w4p, 1w4q, 1w5v, 1w5w, 1w5x, 1w5y, 1w7g, 1w9u, 1w9v, 1w11, 1w13, 1w96, 1wc1, 1wcq, 1wdn, 1wht, 1wm1, 1wn6, 1ws1, 1ws4, 1wuq, 1wur, 1wvj, 1x1z, 1x8d, 1x8j, 1x8r, 1x8t, 1x38, 1x39, 1xap, 1xbo, 1xd0, 1xff, 1xgi, 1xh4, 1xh5, 1xh9, 1xhy, 1xjd, 1xk5, 1xk9, 1xka, 1xkk, 1xow, 1xpz, 1xq0, 1xr9, 1xt8, 1xug, 1xws, 1y0l, 1y1z, 1y3n, 1y3p, 1y3v, 1y3x, 1y6q, 1y20, 1yc4, 1yda, 1ydb, 1ydd, 1ydk, 1yds, 1yei, 1yej, 1yet, 1yfz, 1yp9, 1ype, 1ypg, 1ypj, 1yq7, 1yqj, 1yqy, 1yvm, 1z4o, 1z6s, 1z9y, 1z71, 1zc9, 1zdp, 1zea, 1zfq, 1zge, 1zgi, 1zhy, 1zoe, 1zog, 1zoh, 1zp8, 1zpa, 1zs0, 1zsf, 1zvx, 2a4m, 2a5b, 2a5c, 2a5s, 2a8g, 2a14, 2aac, 2afw, 2afx, 2aj8, 2am4, 2amt, 2ans, 2aoc, 2aod, 2aoe, 2aog, 2aqu, 2arm, 2avm, 2avo, 2avq, 2avs, 2ax9, 2ayr, 2azr, 2b1g, 2b1i, 2b4l, 2b07, 2b7d, 2b9a, 2baj, 2bak, 2bal, 2bes, 2bet, 2bfq, 2bfr, 2bmk, 2bo4, 2boh, 2boj, 2bok, 2bpv, 2bpy, 2bq7, 2bqv, 2brm, 2bt9, 2buv, 2bvd, 2bvr, 2bvs, 2byr, 2bys, 2bz6, 2bza, 2c1p, 2c3l, 2c80, 2c92, 2c94, 2c97, 2ca8, 2cbj, 2cbu, 2cbz, 2cc7, 2ccb, 2ccc, 2ce9, 2cej, 2cen, 2ceq, 2cer, 2ces, 2cex, 2cf8, 2cf9, 2cgf, 2cgr, 2cht, 2cle, 2clh, 2cli, 2clk, 2cn0, 2csn, 2ctc, 2d0k, 2d1n, 2d1o, 2d3u, 2d3z, 2doo, 2drc, 2dri, 2dw7, 2e1w, 2e2p, 2e2r, 2e7f, 2e9u, 2e27, 2e91, 2e92, 2e94, 2epn, 2erz, 2euk, 2evl, 2ewa, 2ewb, 2ews, 2exm, 2ez7, 2f1g, 2f6t, 2f7i, 2f7o, 2f7p, 2f8g, 2f9k, 2f34, 2f35, 2f80, 2f81, 2f94, 2fdp, 2fgu, 2fgv, 2fle, 2flr, 2fmb, 2fpz, 2fqo, 2fqt, 2fqw, 2fqx, 2fqy, 2fu8, 2fw6, 2fx6, 2fxu, 2fxv, 2fzc, 2fzk, 2g5u, 2g94, 2gh9, 2gj5, 2gkl, 2gl0, 2glp, 2gss, 2gst, 2gsu, 2gv6, 2gv7, 2gvj, 2gvv, 2gyi, 2gz2, 2gzl, 2h3e, 2h4g, 2h4k, 2h4n, 2h6b, 2h6t, 2h15, 2h21, 2ha2, 2ha3, 2ha6, 2hah, 2haw, 2hb3, 2hhn, 2hjb, 2hkf, 2hl4, 2hmu, 2hmv, 2hnc, 2hnx, 2hoc, 2hs1, 2hu6, 2hxm, 2hzy, 2i0a, 2i2c, 2i3h, 2i3i, 2i4d, 2i4j, 2i4u, 2i4v, 2i4w, 2i4x, 2i4z, 2i6b, 2i19, 2i80, 2idw, 2ihj, 2ihq, 2iko, 2isw, 2iuz, 2izl, 2j2u, 2j4g, 2j4i, 2j7b, 2j7d, 2j7e, 2j7f, 2j7g, 2j27, 2j34, 2j47, 2j62, 2j75, 2j77, 2j79, 2j94, 2j95, 2jdm, 2jdp, 2jds, 2jdu, 2jew, 2jf4, 2jfz, 2jg0, 2jgs, 2jh0, 2jh5, 2jh6, 2jiw, 2jjb, 2jke, 2jkh, 2jkp, 2jxr, 2mas, 2nmx, 2nmz, 2nn1, 2nn7, 2nnd, 2nsj, 2nt7, 2nta, 2o0u, 2o4j, 2o4k, 2o4l, 2o4n, 2o4r, 2o8h, 2oag, 2oax, 2oc2, 2ogy, 2oi0, 2oi2, 2oiq, 2ojg, 2ojj, 2olb, 2ole, 2on6, 2ovv, 2ovy, 2oxd, 2oxn, 2oxx, 2oxy, 2oym, 2p2a, 2p3a, 2p3b, 2p3c, 2p3i, 2p4j, 2p4s, 2p7a, 2p7g, 2p7z, 2p09, 2p16, 2p53, 2p95, 2pbw, 2pcp, 2pk5, 2pk6, 2pou, 2pov, 2pow, 2pq9, 2pqb, 2pqc, 2pql, 2pqz, 2psu, 2psv, 2ptz, 2pu1, 2pu2, 2pv1, 2pvh, 2pvj, 2pvk, 2pvl, 2pvm, 2pvu, 2pwc, 2pwd, 2pwg, 2pwr, 2py4, 2pym, 2pyn, 2pyy, 2q1q, 2q2a, 2q5k, 2q6f, 2q7q, 2q8m, 2q8z, 2q38, 2q54, 2q55, 2q63, 2q64, 2q88, 2q89, 2qbs, 2qbu, 2qbw, 2qci, 2qd6, 2qd7, 2qd8, 2qdt, 2qg0, 2qg2, 2qhy, 2qhz, 2qi0, 2qi1, 2qi3, 2qi4, 2qi5, 2qi6, 2qi7, 2qm9, 2qmg, 2qnn, 2qnp, 2qpq, 2qpu, 2qrk, 2qrl, 2qta, 2qtg, 2qtn, 2qtt, 2qu6, 2qw1, 2qwb, 2qwc, 2qwd, 2qwe, 2qwf, 2r0h, 2r0z, 2r1y, 2r2m, 2r2w, 2r3t, 2r3w, 2r5a, 2r5p, 2r9x, 2r23, 2r38, 2r43, 2r58, 2r59, 2r75, 2ra0, 2ra6, 2rcb, 2rcn, 2rd6, 2reg, 2rfh, 2ri9, 2rin, 2rio, 2rka, 2rkd, 2rke, 2rkf, 2rkg, 2rkm, 2sim, 2std, 2tmn, 2tpi, 2usn, 2uwd, 2uwl, 2uwo, 2uwp, 2uxi, 2uxz, 2uy0, 2uy3, 2uy4, 2uy5, 2uyn, 2uyq, 2uz9, 2v2c, 2v2h, 2v2q, 2v2v, 2v3d, 2v3u, 2v25, 2v54, 2v57, 2v58, 2v59, 2v77, 2v88, 2v95, 2vb8, 2vba, 2vc9, 2ves, 2vfk, 2vh0, 2vh6, 2vhj, 2vj8, 2vjx, 2vk6, 2vl4, 2vmc, 2vmd, 2vmf, 2vnp, 2vnt, 2vo4, 2vo5, 2vot, 2vpe, 2vpn, 2vpo, 2vqt, 2vrj, 2vsl, 2vt3, 2vuk, 2vvc, 2vvs, 2vvu, 2vvv, 2vw1, 2vw2, 2vwc, 2vwl, 2vwm, 2vwn, 2vwo, 2vxn, 2vyt, 2vzr, 2w5g, 2w08, 2w8j, 2w8w, 2w8y, 2w9h, 2w26, 2w47, 2w67, 2wb5, 2wc3, 2wc4, 2we3, 2web, 2wec, 2wed, 2weh, 2wej, 2weo, 2weq, 2wf5, 2wgj, 2whp, 2wjg, 2wk6, 2wky, 2wkz, 2wl0, 2wly, 2wlz, 2wm0, 2wnj, 2wor, 2wos, 2wq5, 2wr8, 2wuf, 2wvz, 2wyf, 2wyg, 2wyj, 2wzf, 2wzm, 2wzs, 2x0y, 2x2r, 2x4z, 2x6x, 2x7t, 2x8z, 2x09, 2x91, 2x95, 2x96, 2x97, 2xab, 2xb7, 2xbp, 2xbw, 2xbx, 2xc0, 2xc4, 2xd9, 2xda, 2xde, 2xdk, 2xdx, 2xef, 2xeg, 2xei, 2xej, 2xg9, 2xhm, 2xht, 2xib, 2xj1, 2xj2, 2xjg, 2xjj, 2xjx, 2xm1, 2xmy, 2xn3, 2xn5, 2xog, 2xp7, 2xpk, 2xxr, 2xxt, 2xxx, 2xyd, 2xye, 2xyf, 2xyt, 2y5f, 2y5g, 2y7i, 2y7x, 2y7z, 2y8c, 2y80, 2y81, 2y82, 2ya6, 2ya7, 2ya8, 2yay, 2yaz, 2yb0, 2ydt, 2ydw, 2yek, 2yel, 2yfa, 2yfx, 2ygf, 2yhw, 2yi0, 2yi7, 2yix, 2yk1, 2ylc, 2yme, 2ypi, 2ypo, 2yxj, 2yz3, 2z1w, 2z4o, 2z94, 2za0, 2zc9, 2zcs, 2zdk, 2zdl, 2zdm, 2zdn, 2zfp, 2zfs, 2zft, 2zgx, 2zjw, 2zmm, 2zn7, 2zq0, 2zq2, 2zwz, 2zx6, 2zx7, 2zx8, 2zxd, 2zxg, 2zym, 2zz1, 2zz2, 3a1c, 3a1d, 3a1e, 3a2o, 3a5y, 3a6t, 3aaq, 3aas, 3aau, 3acx, 3agl, 3ahn, 3aho, 3ai8, 3aid, 3alt, 3ao2, 3ao5, 3ap4, 3aqt, 3arw, 3arx, 3axz, 3b2q, 3b3s, 3b3x, 3b4f, 3b4p, 3b7j, 3b7r, 3b7u, 3b24, 3b25, 3b26, 3b50, 3b66, 3b67, 3b92, 3bbb, 3bbf, 3be9, 3bex, 3bft, 3bfu, 3bgb, 3bgc, 3bgq, 3bgs, 3bkk, 3bkl, 3bl0, 3bl1, 3bpc, 3bqc, 3bra, 3brn, 3bu1, 3buf, 3bug, 3buh, 3bva, 3bvb, 3bwj, 3bxe, 3bxf, 3bxg, 3bxh, 3bzf, 3c2f, 3c2o, 3c2r, 3c2u, 3c4h, 3c8a, 3c8b, 3c39, 3c52, 3c56, 3c79, 3c84, 3c88, 3c89, 3cct, 3ccw, 3ccz, 3cd0, 3cd5, 3cd7, 3cda, 3cdb, 3cf8, 3cfn, 3cft, 3cj2, 3cj5, 3ckb, 3ckp, 3cm2, 3cow, 3cs7, 3ctt, 3cyw, 3cyx, 3cyz, 3cz1, 3czv, 3d0b, 3d0e, 3d1x, 3d1y, 3d1z, 3d2e, 3d4y, 3d6o, 3d6p, 3d7k, 3d7z, 3d8w, 3d8z, 3d9z, 3d50, 3d51, 3d52, 3d78, 3d83, 3d91, 3da9, 3daz, 3dbu, 3dc3, |

| | |
|---|---|
| | 3dcc, 3dd8, 3ddf, 3ddg, 3dgo, 3djk, 3djo, 3djp, 3djq, 3djv, 3djx, 3dk1, 3dln, 3dnd, 3dne, 3dp4, 3dp9, 3drf, 3drg, 3dri, 3dsz, 3dx3, 3dx4, 3dyo, 3dzt, 3e3c, 3e5u, 3e6y, 3e12, 3eax, 3eb1, 3ebh, 3ebi, 3ebl, 3ebo, 3ed0, 3eeb, 3eft, 3egt, 3ehx, 3ejp, 3ejq, 3eko, 3ekp, 3ekr, 3ekt, 3ekv, 3ekw, 3ekx, 3el1, 3el4, 3el5, 3el9, 3elc, 3eqr, 3ery, 3evd, 3ewc, 3ewj, 3exe, 3exh, 3f1a, 3f5j, 3f5k, 3f5l, 3f6e, 3f6g, 3f7g, 3f7h, 3f7i, 3f8c, 3f8e, 3f8f, 3f15, 3f16, 3f17, 3f18, 3f19, 3f37, 3f48, 3f68, 3f70, 3f78, 3f80, 3fas, 3fat, 3fed, 3fee, 3ff3, 3ffg, 3ffp, 3fh7, 3fhb, 3fj7, 3fjg, 3fl5, 3fqe, 3fql, 3fuc, 3fuz, 3fv3, 3fvh, 3fvk, 3fvl, 3fvn, 3fwv, 3fx6, 3fzn, 3fzy, 3g0e, 3g0i, 3g1d, 3g1v, 3g2y, 3g3r, 3g5k, 3g19, 3g30, 3g32, 3g34, 3g35, 3ga5, 3gba, 3gbe, 3gcp, 3gcs, 3gcu, 3gdt, 3ggu, 3gi4, 3gi5, 3gi6, 3gjw, 3gk1, 3gkz, 3gm0, 3gqz, 3gs6, 3gsm, 3gss, 3gst, 3gt9, 3gta, 3gtc, 3gvb, 3gx0, 3gy2, 3gy3, 3gy7, 3h1x, 3h5b, 3h8b, 3h30, 3h78, 3h89, 3hb4, 3hcm, 3hek, 3hf8, 3hfb, 3hig, 3hit, 3hk1, 3hkn, 3hkq, 3hkt, 3hku, 3hkw, 3hky, 3hl5, 3hl7, 3hl8, 3hll, 3hmo, 3hmp, 3hp9, 3hs4, 3hu3, 3hub, 3huc, 3hv8, 3hvi, 3hvj, 3hww, 3hzk, 3hzm, 3hzv, 3i3b, 3i4y, 3i5z, 3i6o, 3i7e, 3i9g, 3i25, 3i51, 3i60, 3i73, 3iae, 3ibi, 3ibl, 3ibn, 3ibu, 3ies, 3ifl, 3igp, 3ijh, 3ikd, 3ikg, 3imc, 3ime, 3iob, 3ioc, 3iod, 3ioe, 3iof, 3iog, 3ip5, 3ip6, 3ip8, 3ip9, 3iph, 3ipu, 3iqu, 3isj, 3iss, 3iub, 3iue, 3ivc, 3ivx, 3iw5, 3iw6, 3iww, 3jdw, 3jrs, 3jrx, 3juk, 3juo, 3jup, 3jy0, 3jyr, 3jzh, 3jzj, 3k00, 3k02, 3k2f, 3k4d, 3k4q, 3k8c, 3k8o, 3k8q, 3k37, 3k97, 3k99, 3kdb, 3kdc, 3kdd, 3kdm, 3kek, 3kgq, 3kgt, 3kgu, 3kiv, 3kjd, 3kku, 3kmc, 3kmx, 3kmy, 3kqr, 3kr4, 3kv2, 3kyq, 3l0v, 3l3l, 3l3m, 3l3n, 3l4u, 3l4v, 3l4w, 3l4x, 3l4y, 3l4z, 3l59, 3ldp, 3ldq, 3le9, 3lea, 3lgs, 3lir, 3liw, 3ljg, 3ljo, 3ljz, 3lk8, 3lmk, 3lp4, 3lp7, 3lpi, 3lpk, 3lpl, 3lpp, 3lq2, 3lvw, 3lxe, 3lxk, 3lzs, 3lzu, 3lzz, 3m1k, 3m3c, 3m3x, 3m3z, 3m5e, 3m6r, 3m8u, 3m35, 3m36, 3m37, 3m40, 3m67, 3m96, 3mam, 3mdz, 3mf5, 3mfv, 3mfw, 3mhc, 3mhi, 3mhl, 3mhm, 3mho, 3mhw, 3miy, 3mjl, 3ml2, 3ml5, 3mmf, 3mna, 3muz, 3mv0, 3mxd, 3mxe, 3myq, 3mz6, 3mzc, 3n0n, 3n1c, 3n2p, 3n2u, 3n2v, 3n3g, 3n3j, 3n4b, 3n7o, 3n8k, 3n9r, 3n9s, 3n35, 3nb5, 3nee, 3neo, 3nes, 3nex, 3ng4, 3nhi, 3nht, 3ni5, 3nik, 3nim, 3nkk, 3nox, 3npc, 3nq3, 3nsn, 3nu3, 3nu4, 3nu5, 3nu6, 3nu9, 3nuj, 3nuo, 3nw3, 3nxq, 3nyd, 3nyx, 3nzk, 3o4k, 3o5n, 3o5x, 3o7u, 3o8p, 3o9a, 3o9d, 3o9e, 3o9p, 3o56, 3o75, 3o84, 3o99, 3oaf, 3ocp, 3ocz, 3ohi, 3oil, 3oim, 3ok9, 3oku, 3old, 3ouj, 3ov1, 3ove, 3ovn, 3owj, 3own, 3oy0, 3oyq, 3ozg, 3ozj, 3ozp, 3ozr, 3p2e, 3p3g, 3p3r, 3p3s, 3p3t, 3p4v, 3p5l, 3p7i, 3p8n, 3p8o, 3p8p, 3p8z, 3p9l, 3p9m, 3p17, 3p58, 3pb7, 3pb8, 3pb9, 3pbb, 3pce, 3pcf, 3pcg, 3pcj, 3pck, 3pcn, 3pd8, 3pd9, 3pe1, 3pe2, 3pfp, 3pgl, 3pgu, 3pju, 3pn1, 3pn4, 3po1, 3po6, 3ppm, 3ppp, 3ppq, 3ppr, 3ps1, 3pwd, 3pwk, 3pwm, 3q1x, 3q2j, 3q6w, 3q6z, 3q7q, 3q44, 3q71, 3qaa, 3qbc, 3qdd, 3qfd, 3qfy, 3qfz, 3qgw, 3qkd, 3qlm, 3qox, 3qps, 3qqa, 3qt6, 3qto, 3qtv, 3qw5, 3qwc, 3qx5, 3qx9, 3qxt, 3qxv, 3r1v, 3r4m, 3r4n, 3r4p, 3r5t, 3r6u, 3r7o, 3r16, 3r17, 3r24, 3rbu, 3re4, 3rf4, 3rf5, 3rlb, 3rlp, 3rlq, 3rm4, 3rm9, 3roc, 3rt8, 3rtf, 3ru1, 3rux, 3rv4, 3rv8, 3rwp, 3ryv, 3ryx, 3ryy, 3ryz, 3rz0, 3rz1, 3rz5, 3rz7, 3rz8, 3s0b, 3s0d, 3s0e, 3s2v, 3s5y, 3s6t, 3s8l, 3s8n, 3s8o, 3s9e, 3s43, 3s45, 3s54, 3s71, 3s72, 3s73, 3s75, 3s76, 3s77, 3s78, 3sfg, 3sha, 3shc, 3si3, 3si4, 3sio, 3sjf, 3sk2, 3slz, 3sm2, 3spf, 3sr4, 3st5, 3std, 3str, 3su0, 3su1, 3su2, 3su3, 3su4, 3su5, 3su6, 3sue, 3suf, 3sug, 3sur, 3sus, 3sut, 3suu, 3suv, 3suw, 3sv2, 3sw8, 3sww, 3sxf, 3t0x, 3t1a, 3t1m, 3t2q, 3t2w, 3t3c, 3t3u, 3t5u, 3t6b, 3t08, 3t8v, 3t60, 3t64, 3t70, 3t82, 3t83, 3t84, 3t85, 3ta0, 3ta1, 3tao, 3tay, 3tb6, 3tcg, 3td4, 3tf6, 3tfn, 3tfp, 3tfu, 3th9, 3tif, 3tk2, 3tkw, 3tmk, 3ts4, 3tt4, 3ttm, 3ttp, 3tu7, 3tvc, 3tz0, 3tza, 3tzm, 3u5l, 3u6h, 3u6i, 3u8j, 3u8l, 3u10, 3u81, 3u90, 3u92, 3u93, 3ubd, 3ucj, 3udd, 3ug2, 3uil, 3uj9, 3ujc, 3ujd, 3umq, 3uod, 3upk, 3upv, 3usx, 3uu1, 3uug, 3uw4, 3uw5, 3uxd, 3uxk, 3uxl, 3uyr, 3uz5, 3uzj, 3v2n, 3v2p, 3v2q, 3v4t, 3v5p, 3v5t, 3v7x, 3v51, 3v78, 3vbd, 3vd9, 3vdb, 3veh, 3vf5, 3vf7, 3vfa, 3vfb, 3vh9, 3vha, 3vhc, 3vhd, 3vhk, 3vjc, 3vje, 3vtr, 3vvy, 3vw1, 3vw2, 3vx3, 3w5n, 3w07, 3w9k, 3w9r, 3w37, 3wgg, 3wha, 3wjw, 3wmc, 3wtl, 3wtm, 3wtn, 3wto, 3wvm, 3wz6, 3wz7, 3wzn, 3x00, 3zbx, 3zc5, 3zcl, 3zdh, 3zdv, 3zhx, 3zi0, 3zi8, 3zj6, 3zk6, 3zln, 3zlr, 3zm9, 3znr, 3zns, 3zps, 3zpu, 3zq9, 3zqe, 3zsq, 3zsy, 3zt3, 3zv7, 3zxz, 3zyf, 3zyu, 3zze, 4a4q, 4a4v, 4a4w, 4a6b, 4a6c, 4a6l, 4a6s, 4a7i, 4a95, 4ab9, 4aba, 4abb, 4abd, 4abe, 4abf, 4abh, 4acc, 4aci, 4ad2, 4ad3, 4ad6, 4afg, 4ag8, 4agc, 4agl, 4agm, 4ago, 4ahr, 4ahs, 4ahu, 4ai5, 4aia, 4aj4, 4aje, 4aji, 4ajl, 4alx, 4aoi, 4ap7, 4app, 4aqh, 4ara, 4arb, 4asd, 4ase, 4asj, 4att, 4auj, 4av4, 4av5, 4avh, 4avi, 4avj, 4avs, 4ax9, 4axd, 4ayq, 4ayu, 4az5, 4az6, 4azb, 4azc, 4azg, 4azi, 4b0b, 4b1j, 4b2i, 4b2l, 4b3b, 4b3c, 4b3d, 4b5d, 4b5t, 4b5w, 4b6o, 4b6p, 4b6r, 4b6s, 4b7j, 4b7p, 4b7r, 4b8y, 4b9z, 4b32, 4b33, 4b34, 4b35, 4b73, 4bah, 4bak, 4bam, 4ban, 4bao, 4baq, 4bb9, 4bc5, 4bck, 4bcm, 4bcn, 4bco, 4bcp, 4bcs, 4bf1, 4bf6, 4bi6, 4bi7, 4bj8, 4bks, 4bny, 4bqg, 4bqh, 4bqs, 4br3, 4bt3, 4bt4, 4bt5, 4btk, 4bup, 4buq, 4c1t, 4c1u, 4c1y, 4c2v, 4c5d, 4c6u, 4c9x, 4c52, 4ca5, 4ca6, 4ca8, 4cc5, 4cd0, 4cd4, 4cd5, 4ceb, 4cfl, 4cg8, 4cg9, 4cga, 4cgi, 4cj4, 4cjp, 4cjq, 4cjr, 4ck3, 4cl6, 4clj, 4cmo, 4cp5, 4cp7, 4cpr, 4cps, 4cpt, 4cpw, 4cpy, 4cpz, 4cr5, 4crb, 4crf, 4crl, 4cs9, 4csd, 4css, 4cst, 4cu7, 4cu8, 4cwf, 4cwn, 4cwo, 4cwp, 4cwq, 4cwr, 4cws, 4cwt, 4czs, 4d1j, 4d3h, 4d4d, 4d7b, 4d8z, 4daf, 4db7, 4dbm, 4dcs, 4ddm, 4de0, 4de5, 4del, 4der, 4des, 4det, 4deu, 4dew, 4dfg, 4dhl, 4djo, 4djp, 4djq, 4djr, 4dju, 4djw, 4djx, 4djy, 4dko, 4dkp, 4dkq, 4dkr, 4dmw, 4do4, 4do5, 4dq2, 4dst, 4dsu, 4dsy, 4duh, 4dv8, 4dy6, 4dzy, 4e0x, 4e1k, 4e3g, 4e4l, 4e4n, 4e6d, 4e7r, 4e9u, 4e67, 4eb8, 4ef6, 4efk, 4efs, 4egk, 4ehz, 4ei4, 4ej8, 4ejl, 4ek9, 4elf, |

| | |
|---|---|
| | 4elg, 4elh, 4emf, 4emr, 4en4, 4eo6, 4eoh, 4epy, 4er1, 4er2, 4erf, 4etz, 4eu0, 4euo, 4ew2, 4ew3, 4ewn, 4exs, 4ezr, 4ezx, 4ezz, 4f0c, 4f1l, 4f3k, 4f5y, 4f6u, 4f6w, 4f7v, 4f9u, 4f9y, 4f39, 4fai, 4fcq, 4fev, 4few, 4ffs, 4fht, 4fk6, 4fl1, 4fl2, 4flp, 4fm7, 4fm8, 4fnn, 4fp1, 4fs4, 4fsl, 4fxp, 4fxq, 4fys, 4fz3, 4fzj, 4g0p, 4g0q, 4g0y, 4g0z, 4g4p, 4g5f, 4g8m, 4g8n, 4g8v, 4g8y, 4g90, 4g95, 4gah, 4gbd, 4ge1, 4gfo, 4gg7, 4ggz, 4ghi, 4gih, 4gii, 4gj2, 4gj3, 4gkh, 4gki, 4gny, 4gq4, 4gql, 4gqp, 4gqq, 4gqr, 4gr3, 4gr8, 4gu6, 4gu9, 4gue, 4gzp, 4gzt, 4gzw, 4gzx, 4h3f, 4h3g, 4h3j, 4h7q, 4h42, 4h75, 4h81, 4h85, 4ha5, 4hbm, 4hdb, 4hdf, 4hdp, 4heg, 4hf4, 4hfp, 4hj2, 4hla, 4hp0, 4hpi, 4ht0, 4ht2, 4hu1, 4hw3, 4hy1, 4hym, 4hzm, 4i3z, 4i5c, 4i7j, 4i7k, 4i7l, 4i7m, 4i7p, 4i8n, 4i8w, 4i8x, 4i8z, 4i9h, 4i9u, 4i54, 4i71, 4i72, 4i74, 4ibb, 4ibc, 4ibd, 4ibe, 4ibf, 4ibg, 4ibi, 4ibj, 4ibk, 4idn, 4ido, 4ieh, 4igt, 4ih3, 4ih6, 4iic, 4iid, 4iie, 4iif, 4ij1, 4in9, 4io2, 4io3, 4io4, 4io5, 4io6, 4io7, 4ipi, 4ipj, 4ipn, 4ish, 4isi, 4isu, 4itp, 4iue, 4iuo, 4iva, 4iwz, 4j7d, 4j7e, 4j22, 4j44, 4j45, 4j46, 4j47, 4j48, 4j93, 4jal, 4je7, 4je8, 4jfk, 4jfm, 4jh0, 4jkw, 4jn2, 4jne, 4jpx, 4jpy, 4jsa, 4jss, 4jwk, 4jx9, 4jyb, 4jyc, 4jym, 4jyt, 4jz1, 4jzi, 4k0o, 4k0y, 4k3h, 4k3n, 4k4j, 4k5p, 4k6i, 4k7i, 4k7n, 4k7o, 4k9y, 4k55, 4kao, 4kax, 4kb9, 4kcx, 4keq, 4kfq, 4kif, 4kiu, 4km0, 4km2, 4kmz, 4kn0, 4kn1, 4kni, 4knj, 4knm, 4knn, 4ko8, 4kow, 4kp5, 4kp8, 4kqp, 4ks1, 4ks4, 4ksy, 4kwf, 4kwg, 4kwo, 4kx8, 4kxb, 4kxn, 4kyh, 4kyk, 4kz3, 4kz4, 4kz7, 4l2l, 4l4v, 4l4z, 4l6t, 4l9i, 4l19, 4l50, 4l51, 4lar, 4lbu, 4lch, 4leq, 4lhm, 4lhv, 4lj5, 4lj8, 4ljh, 4lk7, 4lkk, 4lko, 4lkq, 4ll3, 4llj, 4llk, 4llp, 4lm0, 4lm1, 4lm2, 4lm3, 4lm4, 4loh, 4loi, 4loo, 4lov, 4loy, 4lps, 4lrr, 4luz, 4lvt, 4lxd, 4lxz, 4ly1, 4ly9, 4lyw, 4lzr, 4m0e, 4m0f, 4m0r, 4m2r, 4m2u, 4m2v, 4m2w, 4m6u, 4m7j, 4m8e, 4m8h, 4m8x, 4m8y, 4m12, 4m13, 4m14, 4mc1, 4mc2, 4mc6, 4mc9, 4mdn, 4mhy, 4mhz, 4mjp, 4mmm, 4mmp, 4mn3, 4mnp, 4mo4, 4mo8, 4mpn, 4mq6, 4mr3, 4mr6, 4mre, 4mrg, 4mrw, 4mrz, 4msa, 4msc, 4msn, 4mss, 4muf, 4mul, 4muv, 4myd, 4n5d, 4n6g, 4n6z, 4n07, 4n7m, 4n7u, 4n8q, 4n9a, 4n9c, 4na9, 4nbk, 4nbl, 4nbn, 4ncn, 4ndu, 4ngm, 4ngn, 4ngp, 4nh7, 4nh8, 4nj9, 4nja, 4nkt, 4nku, 4nl1, 4nnr, 4non, 4np2, 4np3, 4np9, 4nra, 4nuc, 4nue, 4nvp, 4nwc, 4nxu, 4nxv, 4nyf, 4o0a, 4o0b, 4o0x, 4o0y, 4o2c, 4o2p, 4o3c, 4o3f, 4o04, 4o05, 4o6w, 4o07, 4o09, 4o9v, 4o9w, 4o61, 4o97, 4oag, 4oak, 4oc0, 4oc1, 4oc2, 4oc3, 4oc5, 4ocq, 4oct, 4og3, 4og4, 4oiv, 4oks, 4oma, 4omc, 4omj, 4omk, 4or4, 4or6, 4ou3, 4ovf, 4ovg, 4ovh, 4owv, 4ozj, 4p3h, 4p5d, 4p6c, 4p6w, 4p6x, 4p58, 4pb2, 4pee, 4pf5, 4pft, 4pfu, 4pg9, 4phu, 4pin, 4pmm, 4pnu, 4poh, 4poj, 4pop, 4pow, 4pox, 4pp0, 4pp3, 4pp5, 4pqa, 4psb, 4pum, 4pv5, 4pvx, 4pvy, 4pzv, 4q0k, 4q1w, 4q1x, 4q1y, 4q3t, 4q3u, 4q4o, 4q4p, 4q4q, 4q4r, 4q4s, 4q6d, 4q6e, 4q7p, 4q7s, 4q7v, 4q7w, 4q08, 4q8x, 4q8y, 4q09, 4q9o, 4q9y, 4q19, 4q46, 4q81, 4q83, 4q87, 4q90, 4q93, 4q99, 4qb3, 4qdk, 4qem, 4qer, 4qev, 4qew, 4qf7, 4qf8, 4qf9, 4qfl, 4qfn, 4qfo, 4qfp, 4qgd, 4qge, 4qgi, 4qij, 4qj0, 4qjw, 4qjx, 4ql1, 4qlk, 4qll, 4qnb, 4qp2, 4qpd, 4qpl, 4qrh, 4qsu, 4qsv, 4qtl, 4qxo, 4qy3, 4qyy, 4r0a, 4r3w, 4r4c, 4r4i, 4r4o, 4r4t, 4r5a, 4r5b, 4r5t, 4r06, 4r59, 4r73, 4r74, 4r75, 4r76, 4ra1, 4rak, 4rd0, 4rd3, 4rd6, 4rdn, 4re2, 4re4, 4rfc, 4rfd, 4rfr, 4rhx, 4riu, 4riv, 4rj8, 4rn4, 4rpn, 4rpo, 4rqk, 4rqv, 4rr6, 4rra, 4rrf, 4rrg, 4rsk, 4rux, 4ruy, 4ruz, 4rvr, 4rwj, 4rww, 4ryd, 4s1g, 4sga, 4std, 4tim, 4tjz, 4tkb, 4tkh, 4tkj, 4tln, 4tmk, 4tpw, 4tqn, 4trc, 4ts1, 4tt2, 4tte, 4tu4, 4tun, 4ty6, 4tz2, 4u0w, 4u5n, 4u5o, 4u5s, 4u8w, 4u43, 4u54, 4u73, 4ua8, 4uac, 4uc5, 4ucc, 4ufh, 4ufi, 4ufj, 4ufk, 4ufl, 4ufm, 4uin, 4uj1, 4uj2, 4uja, 4ujb, 4uma, 4umb, 4umc, 4und, 4unp, 4uof, 4uoh, 4up5, 4ury, 4urz, 4us3, 4uye, 4uyf, 4v01, 4v24, 4v27, 4w9d, 4w9f, 4w9j, 4w9k, 4w9o, 4w9p, 4w52, 4w97, 4wa9, 4whs, 4wk1, 4wkb, 4wkn, 4wko, 4wkp, 4wn5, 4wop, 4wov, 4wrb, 4wt2, 4x3k, 4x5p, 4x5q, 4x5r, 4x5y, 4x5z, 4x6m, 4x6n, 4x6o, 4x8o, 4x8u, 4x8v, 4x24, 4x48, 4x50, 4xaq, 4xar, 4xas, 4xip, 4xiq, 4xir, 4xit, 4xk9, 4xmb, 4xmr, 4xo8, 4xoc, 4xoe, 4xt2, 4xtv, 4xtw, 4xtx, 4xty, 4xtz, 4xu0, 4xu1, 4xu2, 4xu3, 4xxh, 4xy8, 4xya, 4y0a, 4y2q, 4y3j, 4y3y, 4y4j, 4y5d, 4y8x, 4y59, 4y79, 4yb5, 4ybk, 4yc0, 4yes, 4ygf, 4yha, 4yhm, 4yho, 4yk0, 4ykj, 4ykk, 4ymb, 4ymg, 4ymh, 4yml, 4ymq, 4ymx, 4ynb, 4ynl, 4yo8, 4yrd, 4ysl, 4ytc, 4yth, 4yx4, 4yxi, 4yyt, 4yzu, 4z0k, 4z0q, 4z1e, 4z1j, 4z1k, 4z2b, 4z07, 4z83, 4z84, 4zae, 4zb6, 4zb8, 4zba, 4zbf, 4zbi, 4zcs, 4zeb, 4zec, 4zei, 4zek, 4zgk, 4zip, 4zji, 4zl4, 4zls, 4zme, 4zo5, 4zow, 4zt8, 4zv1, 4zv2, 4zvi, 4zw5, 4zw6, 4zw7, 4zw8, 4zwx, 4zwz, 4zx0, 4zx1, 4zx3, 4zx4, 4zyf, 4zzd, 4zzx, 4zzy, 4zzz, 5a2i, 5a5q, 5a6k, 5a6x, 5a81, 5aa9, 5aan, 5acy, 5ad1, 5afv, 5ahw, 5alb, 5am6, 5am7, 5amd, 5amg, 5aml, 5ant, 5anu, 5anv, 5aoi, 5aoj, 5aol, 5aqz, 5aut, 5ave, 5avf, 5ayt, 5azf, 5b2d, 5b5f, 5b5g, 5b25, 5boj, 5bry, 5bs4, 5btv, 5btx, 5bv3, 5bw4, 5bwc, 5byi, 5c1w, 5c2a, 5c2o, 5c3p, 5c5t, 5c8n, 5cap, 5caq, 5cas, 5cau, 5cbr, 5cbs, 5cc2, 5cep, 5ceq, 5cj6, 5cjf, 5cks, 5cp5, 5cp9, 5cqt, 5cqu, 5cs3, 5cs6, 5cso, 5csp, 5cst, 5ct2, 5cu4, 5cxa, 5cy9, 5czm, 5d0c, 5d0r, 5d1r, 5d2r, 5d3c, 5d3h, 5d3j, 5d3l, 5d3n, 5d3p, 5d3t, 5d3x, 5d6j, 5d21, 5d24, 5d25, 5d26, 5d45, 5d47, 5d48, 5dbm, 5dex, 5dey, 5dfp, 5dgu, 5dgw, 5dh4, 5dhu, 5dit, 5dkn, 5dlx, 5dnu, 5dpx, 5dq8, 5dqc, 5dqe, 5dqf, 5drr, 5dus, 5duw, 5dw2, 5dx4, 5dxt, 5dyo, 5e1s, 5e2k, 5e2l, 5e2o, 5e2p, 5e2r, 5e2s, 5e3a, 5e6o, 5e7n, 5e8f, 5e13, 5e28, 5e73, 5e74, 5e89, 5edb, 5edc, 5edd, 5edl, 5eei, 5eek, 5een, 5ef7, 5efa, 5efc, 5efh, 5efj, 5egm, 5egu, 5eh5, 5eh7, 5eh8, 5ehq, 5ehr, 5ehv, 5ehw, 5ei3, 5eij, 5eis, 5ekm, 5el9, 5elw, 5en3, 5eng, 5ep7, 5epl, 5epn, 5eq1, 5eqe, 5eqp, 5eqy, 5er1, 5er2, 5er4, 5etb, 5etj, 5eu1, 5ev8, 5evb, 5evd, 5evk, 5evz, 5ew0, 5ewa, 5ewk, 5ewy, 5exl, 5exm, 5exn, 5exw, 5ey0, 5ey4, 5eyr, 5f0f, 5f1h, 5f1r, 5f1v, 5f1x, 5f2p, 5f2r, |

| | |
|---|---|
| | 5f2u, 5f5z, 5f08, 5f8y, 5f9b, 5f25, 5f60, 5f61, 5f62, 5f63, 5f74, 5fbi, 5fck, 5fcz, 5fdc, 5fdi, 5fe6, 5fe7, 5fe9, 5fh7, 5fh8, 5fhm, 5fhn, 5fho, 5fl4, 5fl5, 5fl6, 5flo, 5flq, 5fls, 5flt, 5fnc, 5fnd, 5fnf, 5fng, 5fnr, 5fns, 5fnt, 5fnu, 5fog, 5fol, 5fot, 5fou, 5fov, 5fox, 5fpk, 5fs5, 5fsn, 5fso, 5fsx, 5fsy, 5ftg, 5fto, 5fut, 5fyx, 5g1a, 5g1z, 5g2b, 5g2g, 5g4m, 5g4n, 5g4o, 5g5f, 5g5v, 5g5z, 5g17, 5g45, 5g46, 5g57, 5g60, 5g61, 5gj9, 5gja, 5gmh, 5gmn, 5gof, 5gs9, 5gsa, 5h1t, 5h1u, 5h1v, 5h5f, 5h8e, 5h8g, 5h9r, 5h85, 5ha1, 5hbn, 5hbs, 5hct, 5hcv, 5hcy, 5hi7, 5his, 5hjq, 5hrv, 5hrw, 5hrx, 5htl, 5htz, 5hu9, 5hva, 5hvs, 5hvt, 5hwu, 5hwv, 5hz5, 5hz6, 5hz8, 5hz9, 5i1q, 5i2e, 5i2f, 5i3a, 5i3v, 5i3w, 5i3x, 5i3y, 5i7x, 5i7y, 5i8g, 5i9x, 5i9y, 5i9z, 5i29, 5i80, 5i88, 5ia0, 5ia1, 5ia2, 5ia3, 5ia4, 5ia5, 5ie1, 5igm, 5ih9, 5ihh, 5ii2, 5ijr, 5ikb, 5ime, 5ioz, 5ipc, 5ipj, 5irr, 5isz, 5ito, 5itp, 5ivc, 5ive, 5ivv, 5ivy, 5iwg, 5ix0, 5iyy, 5izf, 5izj, 5j0d, 5j1r, 5j1x, 5j2x, 5j3l, 5j6a, 5j6l, 5j6m, 5j6n, 5j7q, 5j7w, 5j8m, 5j8u, 5j8z, 5j9x, 5j20, 5j27, 5j41, 5j64, 5j82, 5j86, 5ja0, 5jfp, 5jfu, 5jg1, 5jgi, 5jgq, 5jhk, 5ji8, 5jop, 5jox, 5jq5, 5js3, 5jsg, 5jsj, 5jsq, 5jss, 5jt9, 5jvi, 5jxn, 5jxq, 5k0h, 5k0m, 5k1d, 5k1f, 5k03, 5k8s, 5k9w, 5ka1, 5ka7, 5ka9, 5kab, 5kad, 5kat, 5kax, 5kbe, 5kby, 5kcb, 5kej, 5kh3, 5khm, 5kly, 5km9, 5kma, 5ko1, 5ko5, 5kqx, 5kqy, 5kr0, 5kr1, 5kr2, 5kva, 5kz0, 5l2s, 5l3a, 5l4i, 5l4j, 5l4m, 5l7e, 5l7g, 5l8a, 5l8c, 5l8y, 5l9g, 5l9i, 5l9l, 5l9o, 5l30, 5laq, 5ld8, 5ldm, 5ldp, 5lif, 5ljq, 5ljt, 5llc, 5lle, 5llg, 5llh, 5lli, 5llo, 5llp, 5lne, 5lny, 5lom, 5lsg, 5lsh, 5lso, 5ltn, 5lud, 5lvd, 5lvl, 5lvq, 5lvr, 5lwd, 5lwm, 5lyn, 5lyr, 5lz4, 5lz5, 5lz7, 5m04, 5m5d, 5m7s, 5m7u, 5m9w, 5m17, 5m23, 5m25, 5m28, 5m77, 5ma7, 5meh, 5mek, 5mes, 5mg2, 5mge, 5mgf, 5mgj, 5mgk, 5mjn, 5mkr, 5mks, 5mlj, 5mme, 5mmg, 5mn1, 5mnn, 5mnr, 5mo8, 5mod, 5mpk, 5mpn, 5mpz, 5mqe, 5mrb, 5mrm, 5mro, 5mrp, 5msb, 5mwh, 5my8, 5mz8, 5n0d, 5n0e, 5n0f, 5n1r, 5n1s, 5n2t, 5n2z, 5n3v, 5n3y, 5n6s, 5n9r, 5n17, 5n18, 5n24, 5n25, 5n31, 5n34, 5n84, 5n93, 5n99, 5nap, 5nau, 5nbw, 5ndf, 5ne5, 5nea, 5neb, 5nee, 5ngz, 5nih, 5njz, 5nk2, 5nk3, 5nk4, 5nk6, 5nk7, 5nk8, 5nk9, 5nkb, 5nkc, 5nkd, 5nkg, 5nkh, 5nki, 5nlk, 5nn6, 5nvv, 5nvw, 5nvx, 5nw0, 5nw1, 5nw2, 5nw7, 5nwe, 5nwi, 5nxg, 5nxi, 5nxo, 5nxp, 5nxw, 5ny1, 5ny3, 5nya, 5nyh, 5nz4, 5nze, 5nzf, 5nzn, 5o1d, 5o1f, 5o1h, 5o2d, 5o4f, 5o5a, 5o07, 5o9o, 5o9p, 5o9r, 5o9y, 5o58, 5o87, 5oa2, 5oa6, 5oaj, 5od1, 5odx, 5oei, 5ogb, 5oh2, 5oh3, 5oh4, 5oh7, 5oh9, 5oha, 5ohy, 5oku, 5om2, 5om3, 5om7, 5oot, 5op4, 5op5, 5oq8, 5oqu, 5org, 5orh, 5orj, 5ork, 5orv, 5orw, 5os2, 5os4, 5os7, 5os8, 5ose, 5osl, 5oss, 5ost, 5ot8, 5ot9, 5ota, 5otc, 5otr, 5otz, 5ouh, 5ov8, 5ovc, 5ovp, 5ovr, 5ovv, 5ovx, 5owl, 5oxk, 5qa8, 5qal, 5qay, 5qqp, 5qtt, 5std, 5sxm, 5sym, 5sz0, 5sz1, 5sz2, 5sz3, 5sz4, 5sz5, 5sz6, 5sz7, 5t7s, 5t8o, 5t8p, 5t9u, 5t9w, 5t9z, 5t19, 5ta2, 5ta4, 5tb6, 5tbe, 5tbm, 5tcj, 5tef, 5tfx, 5th4, 5ti0, 5tkj, 5tkk, 5tmp, 5tp0, 5tpx, 5ttw, 5tuo, 5tuz, 5twj, 5txy, 5ty9, 5tya, 5u0d, 5u0e, 5u0f, 5u0g, 5u0y, 5u0z, 5u4b, 5u4d, 5u6j, 5u8c, 5u11, 5u12, 5u13, 5u14, 5u28, 5u49, 5uc4, 5ucj, 5ueu, 5uez, 5uf0, 5ufc, 5uff, 5ufp, 5ufr, 5ufs, 5uk8, 5ula, 5uln, 5ulp, 5ult, 5umx, 5umy, 5uoo, 5uov, 5upe, 5upf, 5upj, 5upz, 5ut6, 5uv2, 5uxf, 5v0n, 5v7a, 5v79, 5v82, 5var, 5vb5, 5vb6, 5vb7, 5vc3, 5vc4, 5vcv, 5vcw, 5vcy, 5vcz, 5vd0, 5vd1, 5vd2, 5vd3, 5vgy, 5vh0, 5vi6, 5vih, 5vij, 5vja, 5vkc, 5vl2, 5vm0, 5vo1, 5voj, 5vp9, 5vr8, 5vsf, 5vsj, 5vyy, 5w1e, 5w2s, 5w44, 5wa5, 5wa8, 5wa9, 5wal, 5wbm, 5wbo, 5wcm, 5we9, 5wex, 5wgp, 5wij, 5win, 5wl0, 5wlo, 5wmg, 5wp5, 5wqc, 5wuk, 5wxh, 5wyx, 5wyz, 5x54, 5x62, 5x74, 5xg5, 5xmx, 5xo7, 5xpi, 5xsr, 5xva, 5xvg, 5y8y, 5y12, 5y13, 5y94, 5ya5, 5yas, 5yfs, 5yft, 5yj8, 5yjm, 5yl2, 5ymx, 5yqx, 5yum, 5yyf, 5yz2, 5z5f, 5z7b, 5z7j, 5z66, 5z99, 5za7, 5za8, 5za9, 5zae, 5zaf, 5zag, 5zaj, 5zc5, 5zdb, 5zg0, 5zg1, 5zg3, 5zhl, 5zkc, 5znc, 5zni, 5zo8, 5zvw, 5zw6, 5zyg, 5zyl, 5zym, 6a87, 6abx, 6ajv, 6ajy, 6ajz, 6aox, 6aqs, 6arm, 6arn, 6aro, 6as8, 6ayi, 6ayo, 6ayq, 6ayr, 6ays, 6ayt, 6b1k, 6b4d, 6b4l, 6b4n, 6b4u, 6b5q, 6b7a, 6b7b, 6b59, 6b96, 6b97, 6b98, 6bbx, 6bdy, 6bhv, 6bm5, 6bm6, 6bs3, 6bs4, 6c0s, 6c2r, 6c7w, 6c7x, 6c9s, 6c9v, 6c85, 6cbf, 6cbg, 6cdj, 6cdl, 6cdo, 6cdp, 6ce6, 6ced, 6cfc, 6chp, 6cjr, 6cjs, 6cjv, 6ckr, 6cks, 6ckw, 6cn5, 6cpa, 6cq1, 6csp, 6csq, 6csr, 6css, 6cvf, 6cvv, 6cwh, 6cwn, 6czb, 6czc, 6cze, 6czf, 6d1a, 6d1b, 6d1d, 6d1g, 6d1h, 6d1i, 6d1j, 6d1k, 6d2o, 6d3q, 6d5e, 6d5g, 6d5h, 6d5j, 6d9s, 6d9x, 6d15, 6d16, 6d17, 6d18, 6d19, 6d50, 6d55, 6d56, 6d78, 6dai, 6dak, 6dar, 6dd0, 6det, 6dgl, 6dgq, 6dh1, 6dh2, 6dh6, 6dh7, 6dh8, 6dif, 6dil, 6dj1, 6dj2, 6dj5, 6dj7, 6dl2, 6dpt, 6dpx, 6dpy, 6dpz, 6dq4, 6dsp, 6dy7, 6dyn, 6dyr, 6dys, 6dyu, 6dyv, 6dyw, 6dyy, 6dyz, 6dz0, 6dz2, 6dz3, 6dzx, 6e1y, 6e1z, 6e4a, 6e5l, 6e5t, 6e6m, 6e7j, 6e7r, 6e7t, 6e7u, 6e9a, 6e22, 6e23, 6ebe, 6ecz, 6edr, 6eeb, 6eed, 6eeo, 6efj, 6ei5, 6eif, 6eij, 6eiq, 6eis, 6eiz, 6ej3, 6ekq, 6el5, 6eln, 6elo, 6elp, 6en5, 6eog, 6eol, 6ep4, 6epa, 6epy, 6epz, 6eq1, 6eq2, 6eq3, 6eq5, 6eq7, 6eq8, 6eqp, 6equ, 6eqv, 6eqw, 6eqx, 6ets, 6euc, 6euw, 6eux, 6evn, 6evr, 6ex1, 6exi, 6exj, 6exs, 6ey8, 6ey9, 6eya, 6eyb, 6eyt, 6ezq, 6f1j, 6f1n, 6f3b, 6f05, 6f9g, 6f9u, 6f9v, 6f20, 6f28, 6f90, 6f92, 6fa4, 6faa, 6faf, 6fag, 6fba, 6fcj, 6fe0, 6fgg, 6fh3, 6fhk, 6fhq, 6fhu, 6fmc, 6fmj, 6fnf, 6fng, 6fni, 6fnj, 6fnq, 6fnr, 6fo5, 6fs0, 6fs1, 6fsy, 6ftf, 6ftp, 6ftw, 6ftz, 6fuh, 6fui, 6fuj, 6fv4, 6fyz, 6fz4, 6g0z, 6g2b, 6g2c, 6g2e, 6g2f, 6g2o, 6g2r, 6g2s, 6g3a, 6g3q, 6g3v, 6g5l, 6g5u, 6g6t, 6g7a, 6g9u, 6g9x, 6g14, 6g24, 6g25, 6g27, 6g29, 6g34, 6g35, 6g36, 6g37, 6g38, 6g39, 6gdy, 6ge7, 6gf9, 6gfs, 6gfz, 6gg4, 6gga, 6ggb, 6ggf, 6ghh, 6ghj, 6gj6, 6gj8, 6gji, 6gjj, 6gjl, 6gjm, 6gjn, 6gjr, 6gjw, 6gl8, 6gl9, 6gla, 6glb, 6gnm, 6gnp, 6gnr, 6gnw, 6gon, 6got, 6guc, 6gue, 6guh, 6guk, 6gvf, 6gvz, 6gw4, 6gwe, 6gwr, 6gxb, 6gxe, 6gxg, 6gxq, 6gzd, 6gzm, 6h1d, |

| | |
|---|---|
| | 6h1u, 6h2t, 6h2z, 6h5x, 6h8s, 6h29, 6h33, 6h34, 6h36, 6h37, 6h38, 6h77, 6hai, 6hax, 6hay, 6haz, 6hd6, 6hgf, 6hgg, 6hgi, 6hgj, 6hgr, 6hgs, 6hh3, 6hh5, 6hhr, 6hk3, 6hk4, 6hke, 6hlx, 6hly, 6hm1, 6hmg, 6hni, 6hoq, 6hpw, 6hqy, 6hr2, 6hrq, 6hsh, 6ht1, 6htg, 6hza, 6hzb, 6hzy, 6i0z, 6i5g, 6i8m, 6i8t, 6i8y, 6i11, 6i12, 6i13, 6i14, 6i17, 6i18, 6i61, 6i62, 6i63, 6i64, 6i65, 6i66, 6i67, 6ibk, 6ic2, 6idb, 6idg, 6iez, 6if0, 6ift, 6iht, 6iiu, 6im4, 6ior, 6ios, 6iou, 6ixd, 6j0g, 6j0o, 6j1l, 6j3o, 6j3p, 6j7e, 6j9w, 6j9y, 6j72, 6jad, 6jag, 6jam, 6jan, 6jao, 6jap, 6jaq, 6jav, 6jaw, 6jay, 6jb0, 6jb4, 6jbb, 6jbe, 6jdi, 6jdl, 6jfk, 6jof, 6jon, 6jtc, 6k3l, 6k04, 6kjd, 6m8q, 6ma2, 6ma3, 6ma4, 6ma5, 6md0, 6md6, 6md8, 6mdu, 6mg5, 6mh1, 6mh7, 6mj4, 6mj7, 6mja, 6mjf, 6mji, 6mku, 6mkw, 6ml9, 6mla, 6mle, 6mlg, 6mli, 6mlj, 6mll, 6mln, 6mlo, 6mlp, 6mm2, 6mnc, 6mnv, 6mqc, 6mqe, 6msy, 6mu3, 6mub, 6mxc, 6n0j, 6n0k, 6n3v, 6n3w, 6n3x, 6n3y, 6n5x, 6n7a, 6n7b, 6n7d, 6n8x, 6n9l, 6n78, 6n79, 6n95, 6ncn, 6nco, 6ndl, 6ne5, 6nfh, 6nfy, 6nmb, 6no9, 6np2, 6np3, 6np4, 6np5, 6nsv, 6nu1, 6nu5, 6nv7, 6nv9, 6nw3, 6nwk, 6nwl, 6nxz, 6ny0, 6nyv, 6nzv, 6o0k, 6o0m, 6o0o, 6o0p, 6o5a, 6o5g, 6o5t, 6o5u, 6o5x, 6o9c, 6o48, 6o94, 6o95, 6odz, 6oe0, 6oe1, 6of5, 6oja, 6olx, 6om8, 6ooy, 6oqb, 6oqc, 6oqd, 6orr, 6otg, 6p3t, 6p3v, 6p5o, 6p8a, 6p9e, 6p83, 6p84, 6p85, 6p86, 6p87, 6p88, 6p89, 6pg3, 6pg4, 6pg5, 6pg6, 6pg7, 6pg8, 6pg9, 6pga, 6pgb, 6pgc, 6pge, 6phx, 6pi5, 6pi6, 6pia, 6pid, 6pl1, 6plf, 6poi, 6ppy, 6prf, 6pu3, 6pve, 6pvv, 6pvw, 6pvz, 6py0, 6q3q, 6q3y, 6q3z, 6q4e, 6q4g, 6q54, 6q60, 6qas, 6qau, 6qav, 6qe0, 6qe5, 6qge, 6qgf, 6qgg, 6qgh, 6qi4, 6qi7, 6ql1, 6ql2, 6qln, 6qlo, 6qlp, 6qlq, 6qlr, 6qls, 6qlt, 6qlu, 6qmj, 6qmk, 6qpl, 6qqq, 6qqu, 6qqv, 6qqw, 6qqz, 6qr0, 6qr1, 6qr2, 6qr3, 6qr4, 6qr7, 6qr9, 6qrc, 6quv, 6qz5, 6r0v, 6r1a, 6r1b, 6r1d, 6r1w, 6r4k, 6r8l, 6r8o, 6r8w, 6r9u, 6r9x, 6r11, 6r13, 6rfn, 6rfw, 6rih, 6rnt, 6rot, 6rtw, 6s4n, 6s4t, 6s5k, 6s56, 6s57, 6sbt, 6sge, 6ssy, 6st0, 6std, 6stl, 6szp, 6t1i, 6t1j, 6t1l, 6t1m, 6t1n, 6t1o, 6u5y, 6u6w, 6u8b, 6u8o, 6ud2, 6udi, 6udt, 6udu, 6udv, 6ueg, 6ugp, 6ugr, 6ugz, 6uh0, 6upj, 7std, 7upj, 8a3h, 8cpa, 10gs, 184l, 185l, 186l, 187l, 188l, 456c, 966c |
| PDBbind 2016 core set (Test set) | 1a30, 1bcu, 1bzc, 1c5z, 1e66, 1eby, 1g2k, 1gpk, 1gpn, 1h22, 1h23, 1k1i, 1lpg, 1mq6, 1nc1, 1nc3, 1nvq, 1o0h, 1o3f, 1o5b, 1owh, 1oyt, 1p1n, 1p1q, 1ps3, 1pxn, 1q8t, 1q8u, 1qf1, 1qkt, 1r5y, 1s38, 1sqa, 1syi, 1u1b, 1uto, 1vso, 1w4o, 1y6r, 1yc1, 1ydr, 1ydt, 1z6e, 1z9g, 1z95, 2al5, 2br1, 2brb, 2c3i, 2cbv, 2cet, 2fvd, 2fxs, 2hb1, 2iwx, 2j7h, 2j78, 2p4y, 2p15, 2pog, 2qbp, 2qbq, 2qbr, 2qe4, 2qnq, 2r9w, 2v00, 2v7a, 2vkm, 2vvn, 2vw5, 2w4x, 2w66, 2wbg, 2wca, 2weg, 2wer, 2wn9, 2wnc, 2wtv, 2wvt, 2x00, 2xb8, 2xbv, 2xdl, 2xii, 2xj7, 2xnb, 2xys, 2y5h, 2yfe, 2yge, 2yki, 2ymd, 2zb1, 2zcq, 2zcr, 2zda, 2zy1, 3acw, 3ag9, 3ao4, 3arp, 3arq, 3aru, 3arv, 3ary, 3b1m, 3b5r, 3b27, 3b65, 3b68, 3bgz, 3bv9, 3cj4, 3coy, 3coz, 3d4z, 3d6q, 3dd0, 3dx1, 3dx2, 3dxg, 3e5a, 3e92, 3e93, 3ebp, 3ehy, 3ejr, 3f3a, 3f3c, 3f3d, 3f3e, 3fcq, 3fur, 3fv1, 3fv2, 3g0w, 3g2n, 3g2z, 3g31, 3gbb, 3gc5, 3ge7, 3gnw, 3gr2, 3gv9, 3gy4, 3ivg, 3jvr, 3jvs, 3jya, 3k5v, 3kgp, 3kr8, 3kwa, 3l7b, 3lka, 3mss, 3myg, 3n7a, 3n76, 3n86, 3nq9, 3nw9, 3nx7, 3o9i, 3oe4, 3oe5, 3ozs, 3ozt, 3p5o, 3prs, 3pww, 3pxf, 3pyy, 3qgy, 3qqs, 3r88, 3rlr, 3rr4, 3rsx, 3ryj, 3syr, 3tsk, 3twp, 3u5j, 3u8k, 3u8n, 3u9q, 3udh, 3ueu, 3uev, 3uew, 3uex, 3ui7, 3uo4, 3up2, 3uri, 3utu, 3uuo, 3wtj, 3wz8, 3zdg, 3zso, 3zsx, 3zt2, 4abg, 4agn, 4agp, 4agq, 4bkt, 4cig, 4ciw, 4cr9, 4cra, 4crc, 4ddh, 4ddk, 4de1, 4de2, 4de3, 4djv, 4dld, 4dli, 4e5w, 4e6q, 4ea2, 4eky, 4eo8, 4eor, 4f2w, 4f3c, 4f09, 4f9w, 4gfm, 4gid, 4gkm, 4gr0, 4hge, 4ih5, 4ih7, 4ivb, 4ivc, 4ivd, 4j3l, 4j21, 4j28, 4jfs, 4jia, 4jsz, 4jxs, 4k18, 4k77, 4kz6, 4kzq, 4kzu, 4llx, 4lzs, 4m0y, 4m0z, 4mgd, 4mme, 4ogj, 4owm, 4pcs, 4qac, 4qd6, 4rfm, 4tmn, 4twp, 4ty7, 4u4s, 4w9c, 4w9h, 4w9i, 4w9l, 4wiv, 4x6p, 5a7b, 5aba, 5c2h, 5c28, 5dwr, 5tmn |
| CSAR NRC-HiQ Set1 (Test set) | 1uto, 2add, 1iup, 2zlz, 2cjp, 1ukb, 1tog, 2ou0, 2are, 1ax1, 1w6o, 3f4j, 2jff, 2otz, 2pzv, 3b3c, 1toi, 2ihk, 2qmj, 2jdy, 2arb, 2v8q, 2nsl, 1vso, 2j4k, 2hr6, 2z8f, 2vhw, 2bbf, 2q3c, 2ilz, 1gja, 3d2r, 2jgb, 3c7i, 2v8y, 2pgz, 2v7u, 2b3f, 2qnq, 2p98, 3ene, 3bgz, 2q6m, 2qbr, 2qvu, 2hzl, 1vot, 3f3d, 2v7v, 2rde, 2qbq, 2v7t, 2rca, 2cem, 2vw5, 2qeh, 2qry, 2nnq, 2vkm, 2p4y, 2z4b, 2r6w, 2idz, 2pog, 2jj3, 2jbj, 2p3t |
| CSAR NRC-HiQ Set2 (Test set) | 1rdi, 1q6g, 1y93, 1ps3, 1tok, 1q6e, 2dm5, 1toj, 1gzc, 1gz9, 1bcj, 2hb1, 1bky, 1z3t, 1x9d, 1ax2, 1gww, 1gx0, 2a3c, 1w2g, 1yxd, 1xge, 2brb, 1nc3, 1s38, 2f5t, 1uzv, 1gjd, 1uld, 1gpk, 2hj4, 1syi, 1a8i, 1ha2, 4ubp, 1xw6, 1ow4, 1urg, 1s50, 1s7y, 1gi9, 1nc1, 1yc1, 1hnn, 2fai, 2j78, 1r5y, 1q4w, 1s9t, 2iwx, 1p1n, 1tr7, 1gj8, 2il2, 1z95, 1tt1, 1txf, 1zhx, 1sw1, 1xl5, 2hd6, 1b6j, 1g2k, 1lhw, 2ff1, 2cji, 1b6l, 1h23, 1b6m, 2fvd, 1q0y, 1qkt, 1h22, 1eby, 2fv5 |

**Table S2.** Full list of molecular and binding pocket descriptors used in the study.

| | |
|---|---|
| Molecular descriptors | MolLogP, MolMR, ExactMolWt, HeavyAtomCount, NumHAcceptors, NumHDonors, NumHeteroatoms, NumRotatableBonds, NumAromaticRings, NumAliphaticRings, RingCount, TPSA, LabuteASA, Kappa1, Kappa2, Kappa3, Chi0, Chi1, Chi0n, Chi1n, Chi2n, Chi3n, Chi4n, Chi0v, Chi1v, Chi2v, Chi3v, Chi4v, PEOE_VSA1, PEOE_VSA2, PEOE_VSA3, PEOE_VSA4, PEOE_VSA5, PEOE_VSA6, PEOE_VSA7, PEOE_VSA8, PEOE_VSA9, PEOE_VSA10, PEOE_VSA11, PEOE_VSA12, PEOE_VSA13, PEOE_VSA14, SMR_VSA1, SMR_VSA3, SMR_VSA4, SMR_VSA5, SMR_VSA6, SMR_VSA7, SMR_VSA9, SMR_VSA10, SlogP_VSA1, SlogP_VSA2, SlogP_VSA3, SlogP_VSA4, SlogP_VSA5, SlogP_VSA6, SlogP_VSA7, SlogP_VSA8, SlogP_VSA10, SlogP_VSA11, SlogP_VSA12, EState_VSA1, EState_VSA2, EState_VSA3, EState_VSA4, EState_VSA5, EState_VSA6, EState_VSA7, EState_VSA8, EState_VSA9, EState_VSA10, VSA_EState1, VSA_EState2, VSA_EState3, VSA_EState4, VSA_EState5, VSA_EState6, VSA_EState7, VSA_EState8, VSA_EState9, VSA_EState10 |
| Binding pocket descriptors | pock_vol, nb_AS, mean_as_ray, mean_as_solv_acc, apol_as_prop, mean_loc_hyd_dens, hydrophobicity_score, volume_score, polarity_score, charge_score, flex, prop_polar_atm, as_density, as_max_dst, convex_hull_volume, surf_pol_vdw14, surf_pol_vdw22, surf_apol_vdw14, surf_apol_vdw22, n_abpa, ALA, ARG, ASN, ASP, CYS, GLN, GLU, GLY, HIS, ILE, LEU, LYS, MET, PHE, PRO, SER, THR, TRP, TYR, VAL |

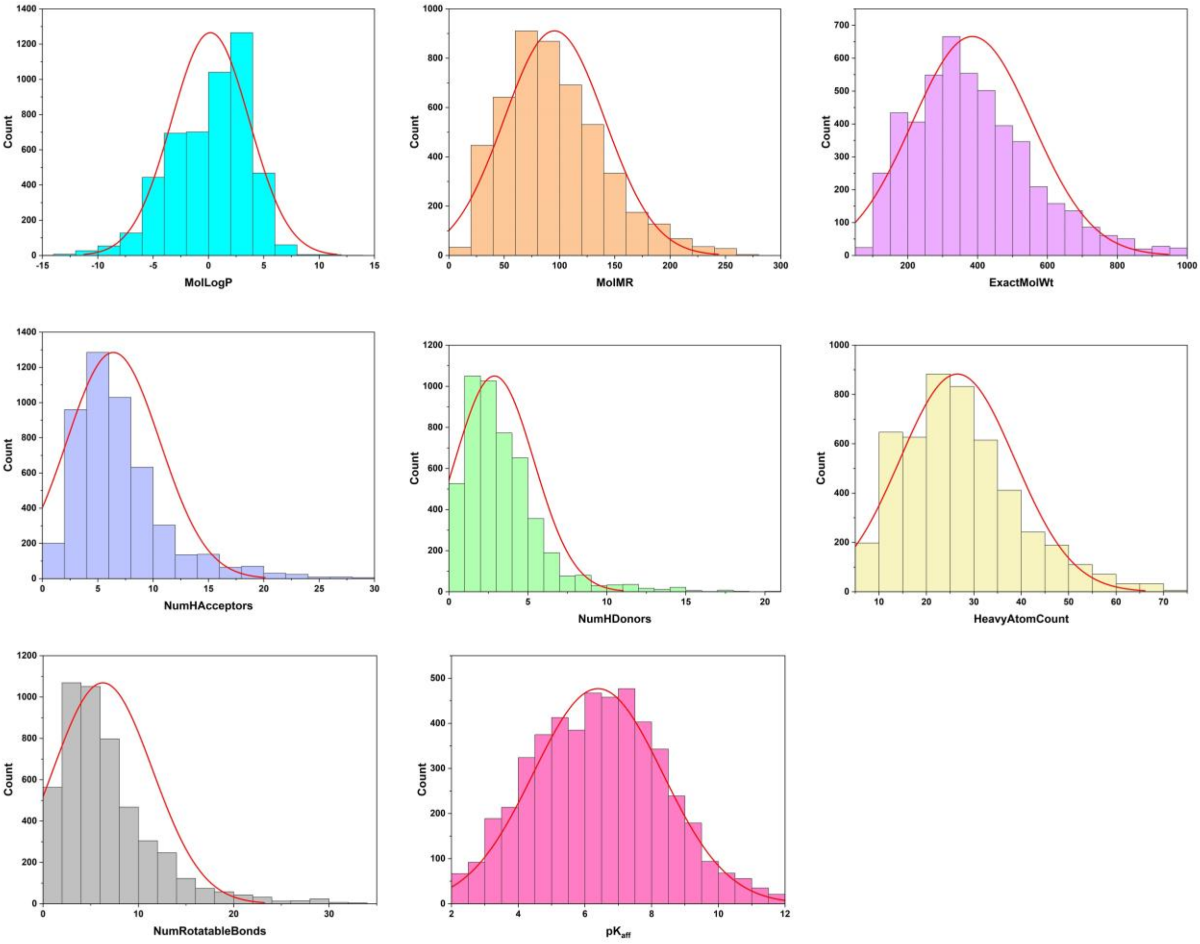


**Figure S1.** Histogram distribution of some molecular descriptors of the ligands in the training set against the logarithmic form of the experimental binding affinity values (*pK<sub>aff</sub>*).

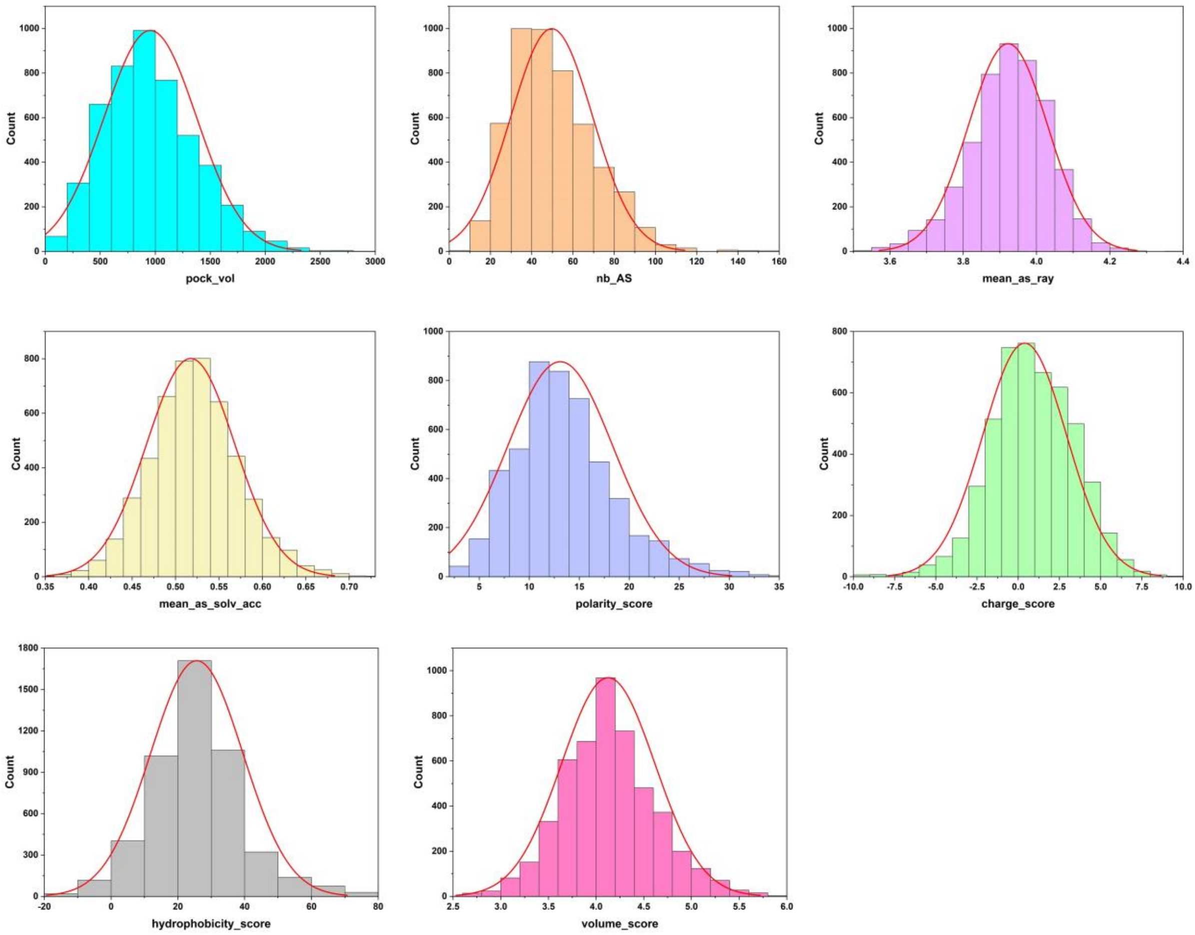


**Figure S2.** Histogram distribution of some binding pocket descriptors in the training dataset.

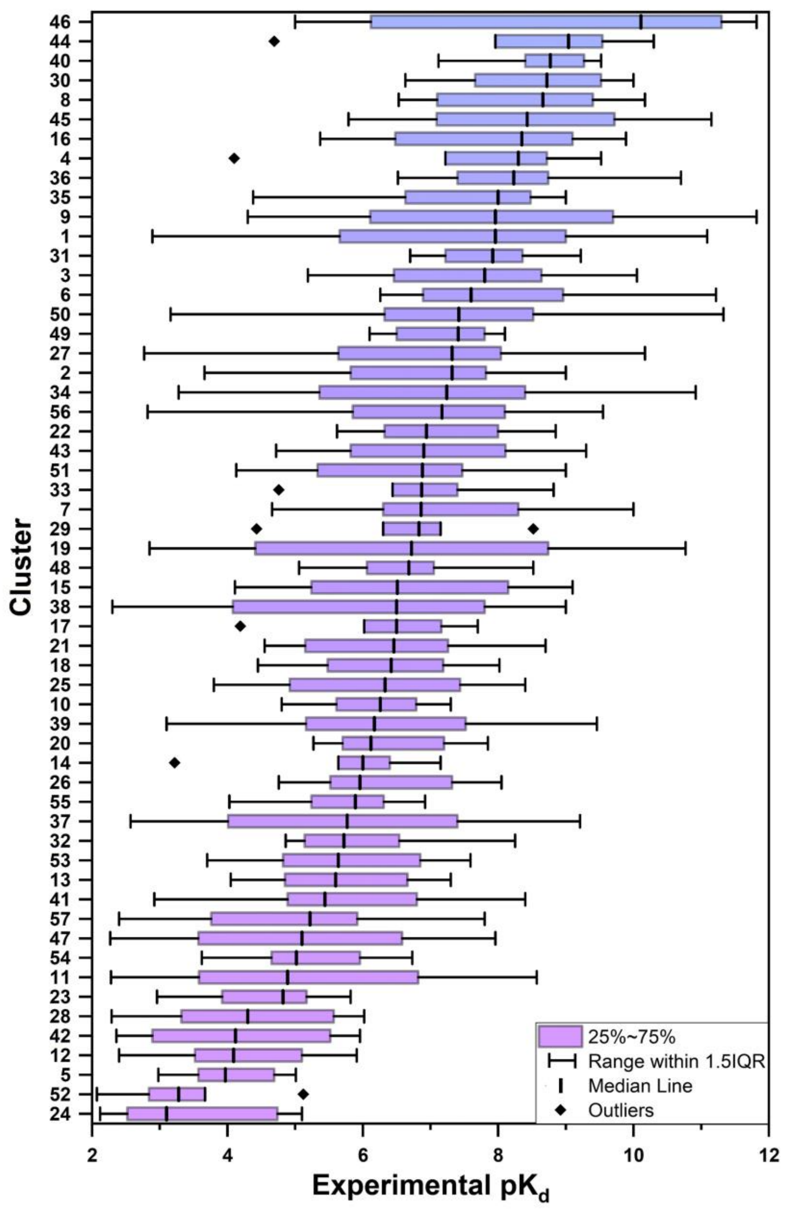


**Figure S3.** Boxplot of experimental binding affinity values for each of the 57 clusters in the PDBbind 2016 core set.
